# Chemogenetic Control of Nanobodies

**DOI:** 10.1101/683557

**Authors:** Helen Farrants, Miroslaw Tarnawski, Thorsten G. Müller, Shotaro Otsuka, Julien Hiblot, Birgit Koch, Moritz Kueblbeck, Hans-Georg Kräusslich, Jan Ellenberg, Kai Johnsson

## Abstract

We introduce an engineered nanobody whose affinity to green fluorescent protein (GFP) can be switched on and off with small molecules. By controlling the cellular localization of GFP fusion proteins, the engineered nanobody allows to study their role in basic biological processes, an approach that should be applicable to numerous previously described GFP fusions. We also outline how the binding affinities of other nanobodies can be controlled by small molecules.

## MAIN TEXT

The variable domains of heavy chain-only antibodies^1^, commonly abbreviated as nanobodies, are powerful tools to interrogate processes in living systems. Nanobodies can be selected to bind to a variety of targets with high affinity and selectivity, and can be functionally expressed inside cells^2,3^. The range of applications of nanobodies would be greatly expanded if their binding affinity towards their target could be rapidly switched on and off with a cell-permeable and non-toxic molecule. Proteins such as kinases and Cas9 have been engineered to control their activity with small molecules^4-6^, but these approaches have not been applied to nanobodies. Here, we introduce “ligand-modulated antibody fragments” (LAMAs), which combine the high selectivity and specificity of nanobodies with the fast temporal control offered through the use of small molecules. LAMAs are generated by inserting a circularly permutated bacterial dihydrofolate reductase (cpDHFR)^7^ into nanobodies. The new termini of this cpDHFR are located in an active site loop of wild-type DHFR. Furthermore, cpDHFR is partially unfolded in the absence of its cofactor nicotinamide adenine dinucleotide phosphate (NADPH) and DHFR inhibitors such as trimethoprim (TMP)^8,9^. TMP is a clinically approved anti-bacterial drug that has excellent cell and tissue permeability and is not toxic for mammalian cells. LAMAs disrupt the binding of the nanobody to its target by exploiting the change in conformation of cpDHFR upon binding of NADPH and DHFR inhibitors (**Fig. 1a**). The first nanobody we subjected to this approach was the enhancer nanobody for GFP^10^. Specifically, we inserted cpDHFR into various sites of the enhancer nanobody and measured the binding affinities of the protein chimeras to wild-type GFP (wtGFP) in the presence and absence of the ligands NADPH and TMP (**Fig. 1b** and **Supplementary Fig. 1**). The most promising insertion hits were in the complementary-determining region 3 (CDR3), which is often essential for making high affinity contacts between nanobodies and their targets^11^. Of these hits, we analyzed ^GFP^LAMA_F98_, ^GFP^LAMA_G97_, and ^GFP^LAMA_N95_ in greater detail (**Fig. 1c-e** and **Supplementary Fig. 2**). All three ^GFP^LAMAs retained a single-digit nanomolar affinity to GFP in the absence of ligands. For all three nanobodies, the affinity towards GFP was dramatically decreased in the presence of NADPH and TMP such that no binding to GFP could be detected for ^GFP^LAMA_F98_ and ^GFP^LAMA_G97_ (**Fig. 1c-e** and **Supplementary Fig. 3**). For ^GFP^LAMA_F98_ the presence of NADPH alone also affected the binding affinity to GFP, whereas the affinity of ^GFP^LAMA_G97_ and ^GFP^LAMA_N95_ was not affected by NADPH.

**Figure 1.**
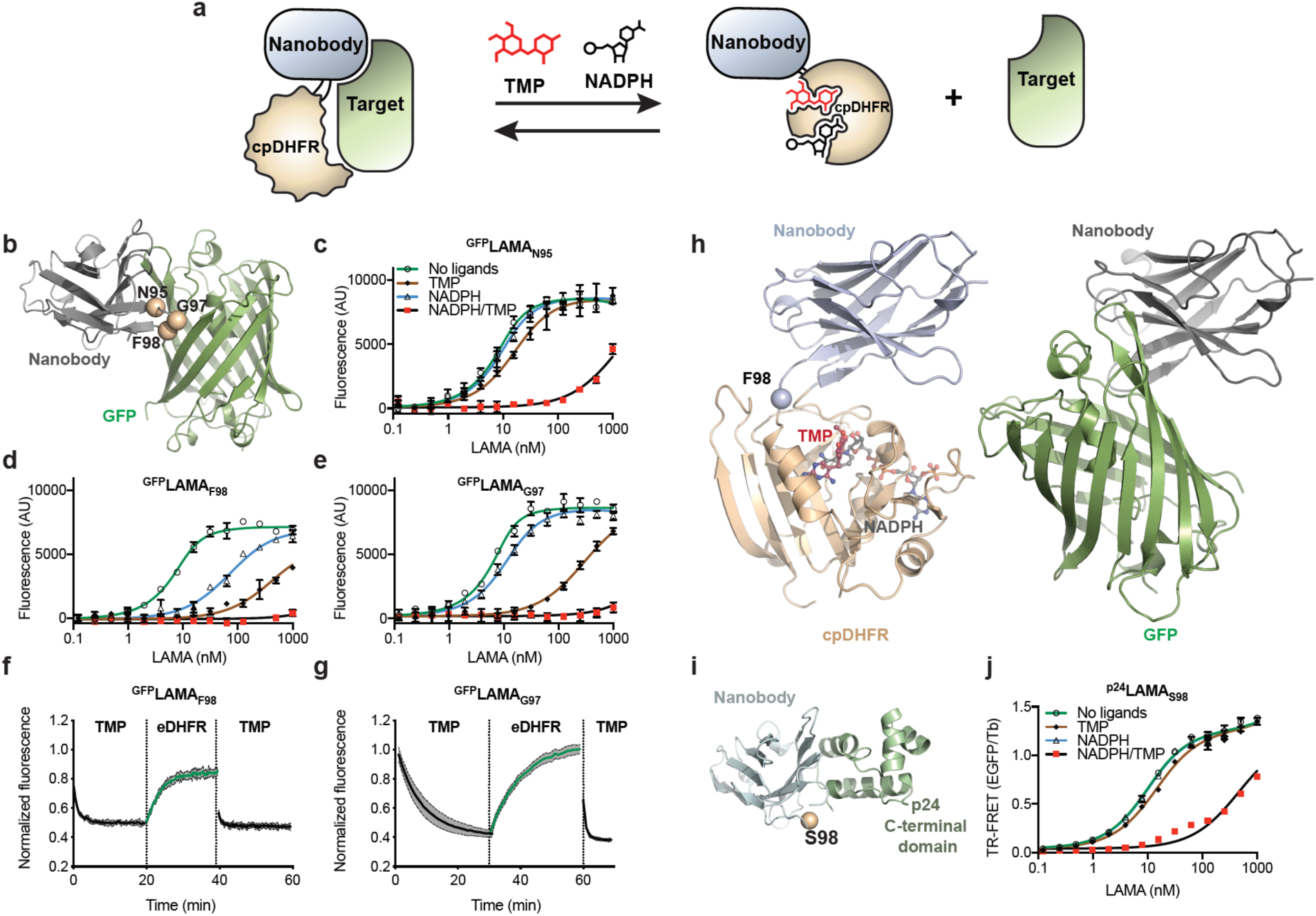
Generation of LAMAs from nanobodies and cpDHFR. (**a**) Schematic illustration of the design principle of LAMAs. (**b**) LAMA insertion positions of cpDHFR highlighted as beige spheres mapped onto the structure of the enhancer nanobody bound to GFP, (PDBID = 3K1K). (**c-e**) Modulation of wtGFP fluorescent emission by ^GFP^LAMA_F98_ (**c**), ^GFP^LAMA_G97_ (**d**), ^GFP^LAMA_N95_ (**e**), in the presence of NADPH (100 *µ*M) and/or TMP (500 *µ*M). Mean ± s.d.. (**f,g**) Dissociation kinetics in the presence of NADPH (100 *µ*M) of ^GFP^LAMA_F98_ (**f**), ^GFP^LAMA_G97_ (**g**), from wtGFP measured by wtGFP emission, on the addition of TMP (1 *µ*M), followed by the competitive removal by eDHFR (8 *µ*M) and addition of excess TMP (50 *µ*M). Mean (solid line) + s.d. (grey area). (**h**) Comparison of the X-ray structure of ^GFP^LAMA_F98_ in the presence of NADPH and TMP (PDBID = 6RUL), with the enhancer nanobody bound to GFP (PDBID = 3K1K). The insertion site F98 is highlighted as a blue sphere. (**i**) LAMA insertion position of cpDHFR highlighted as a beige sphere mapped onto the structure of the nanobody for p24, (PDBID = 2XV6). (**j**) Titration of EGFP-^p24^LAMA_S98_ against Tb-labeled-p24 in a TR-FRET assay in the presence of NADPH (100 *µ*M) and/or TMP (500 *µ*M). Mean ± s.d..

The kinetics of dissociation of the complexes between GFP and ^GFP^LAMA_F98_ or ^GFP^LAMA_G97_ upon addition of TMP were on the timescale of minutes: *t*_1/2_ = 34 ± 1 sec and *t*_1/2_ = 5.6 ± 0.5 min for ^GFP^LAMA_F98_ and ^GFP^LAMA_G97_, respectively (**Fig. 1f,g**). Subsequent removal of TMP by addition of wild-type DHFR resulted in reformation of the complexes within minutes (**Fig. 1f,g**). The complex could then be dissociated again by addition of an excess amount of TMP (**Fig. 1f,g**). The dissociation kinetics of the complexes could also be tuned using DHFR inhibitors with different affinities to DHFR (**Supplementary Fig. 4**). These experiments underline that ^GFP^LAMA_F98_ and ^GFP^LAMA_G97_ can be repeatedly switched on and off through the addition of DHFR inhibitors.

To understand how the cpDHFR insertion into nanobodies allowed control of binding affinities, we solved the crystal structures of ^GFP^LAMA_F98_ and ^GFP^LAMA_G97_ in complex with NADPH and TMP. No major structural changes were seen in the nanobody domain of the two LAMAs relative to enhancer nanobody. Comparing these structures with the structure of enhancer nanobody bound to GFP suggests that folded cpDHFR sterically hampers binding to GFP (**Fig. 1h** and **Supplementary Fig. 5**). The TMP-dependent control of the ^GFP^LAMAs was abolished when GGS-linkers were inserted between cpDHFR and the nanobody (**Supplementary Fig. 6**), indicating that the switching of nanobody affinity did not solely arise from insertion of the protein domain.

Given the large number of nanobodies that have been selected and characterised^12^, we attempted to expand the LAMA concept to other targets. Nanobodies for G-associated kinase^13^, and for lamina-associated polypeptide 1^14^, did not allow for cpDHFR insertion into the tried positions (**Supplementary Fig. 7a,b**). The minimizer nanobody for GFP^10^ greatly decreased its affinity to GFP on cpDHFR insertion, but responded to the addition of ligands (**Supplementary Fig. 7c**). A nanobody for the C-terminal region of the p24 HIV capsid protein (*manuscript in preparation*) could be readily converted into a LAMA on insertion of cpDHFR into the CDR3 loop (**Fig. 1i** and **Supplementary Fig. 8**). The ^p24^LAMA_S98_ showed low nanomolar affinity for p24 HIV capsid protein when no ligands were present. Neither TMP nor NADPH alone could decrease the affinity of ^p24^LAMA_S98_ for its target, but addition of both ligands reduced the affinity 70-fold (**Fig. 1j**). These experiments highlight the transferability for the LAMA approach to other nanobodies.

The binding of both the ^p24^LAMA and ^GFP^LAMAs to their targets could be switched on and off through the addition of TMP in live cells (**Fig 2**). Intracellular NADPH concentration in live cells is estimated to be 3.1 ± 0.3 *µ*M^15^, thus providing a basal level of NADPH. The expression of the cytosolic p24 precursor polyprotein Gag in HIV transfected cells stably expressing an EGFP-^p24^LAMA_S98_ fusion resulted in sequestering of the LAMA in the cytosol (**Fig. 2a**). However, the LAMA was released from the p24 domain of Gag by the addition of TMP within minutes, as demonstrated by diffusion of EGFP-^p24^LAMA_S98_ into the nucleus (**Fig. 2a** and **Supplementary Fig. 9**). Targeting ^GFP^LAMAs to the inner leaflet of the plasma membrane by a Lyn kinase derived sequence^16^ (Lyn-^GFP^LAMA) resulted in localization of EGFP to the plasma membrane, which could be released into the cytosol through addition of TMP (**Fig. 2b** and **Supplementary Fig. 10a**). Similarly, targeting ^GFP^LAMAs to the outer membrane of mitochondria^17^ (mito-^GFP^LAMAs) resulted in reversible sequestering of EGFP to the outer membrane of mitochondria (**Fig. 2c** and **Supplementary Fig. 10b**). The kinetics of the TMP-dependent release and sequestering of EGFP from the outer mitochondrial membrane was evaluated by following the appearance and disappearance of the fluorescence of nuclear EGFP (**Fig. 2d,e** and **Supplementary Fig. 11**). The release and sequestering of EGFP upon addition and wash-out of TMP occurred on a timescale of minutes, and could be repeated over several cycles (**Fig. 2e**). Furthermore, TMP-dependent release of EGFP from mito-^GFP^LAMA_F98_ was dose-dependent up to 5 *µ*M TMP (**Supplementary Fig. 11b**).

**Figure 2.**
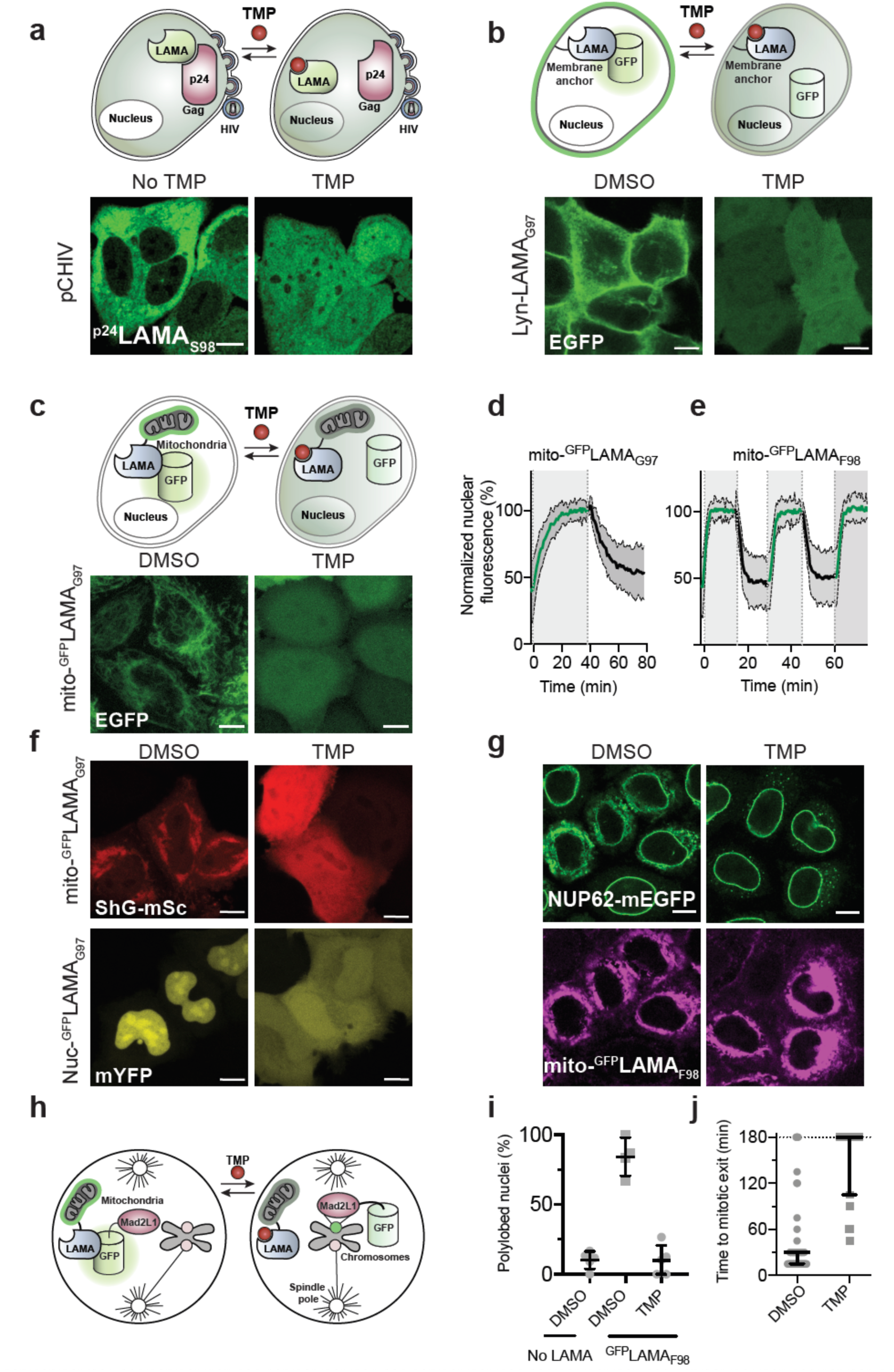
Sequestering and release of protein localization in live cells using LAMAs. (**a**) Schematic illustration and live-cell imaging of sequestering and release of EGFP-^p24^LAMA_S98_ in HeLa TZM-bl cells expressing p24 as part of the Gag polyprotein, after transfection with pCHIV. TMP was added to the cells and followed for 40 minutes. (**b,c**) Schematic illustration and live-cell imaging of HeLa Kyoto cells coexpressing Lyn-^GFP^LAMA_G97_ (**b**) or mito-^GFP^LAMA_G97_ (**c**) with EGFP. Cells were either incubated with DMSO or TMP before imaging. (**d,e**) Kinetics of release and sequestering of EGFP from mito-^GPF^LAMAs in U-2 OS cells. EGFP fluorescence in the nucleus was quantified in cells coexpressing EGFP and (**d**) mito-^GPF^LAMA_G97_ (N = 26 cells) or (**e**) mito-^GPF^LAMA_F98_ (N = 26 cells). Mean (solid line) ± s.d. (grey area). TMP was present in greyed areas. (**f**) Sequestering and release of ShadowG-mScarlet using mito-^GFP^LAMA_G97_ and mYFP using nuc-^GFP^LAMA_G97_ in HeLa Kyoto cells incubated in DMSO or TMP before imaging. (**g**) Genome edited NUP62-mEGFP HeLa cells transiently expressing mito-^GPF^LAMA_F98_ labeled by BG-TMR via a SNAP-tag fusion. Cells were treated with DMSO or TMP before imaging. (**h**) Schematic illustration of Mad2L1-EGFP sequestered away from the centromere to the mitochondria by a ^GFP^LAMA.(**i, j**) Nuclear morphology (**i**) and duration of mitotic events (**j**) during live-cell imaging of Mad2L1-EGFP cells stably expressing mito-^GPF^LAMA_F98,_ after the wash out of TMP (50 *µ*M). Percentage polylobed cells (mean ± s.d., N = 5 independent experiments). Duration of mitotic events recorded in the presence of mitotic arrest drug nocodazole (330 nM) (median ± interquartile range, N = 63 and 19 cells, from 3 independent experiments). TMP = 10 *µ*M, unless otherwise stated. Scale bars, 10 *µ*m.

^GFP^LAMAs can be used to control the localization of other family members of GFP-based proteins to which the enhancer nanobody binds, for example YFP and Shadow G^18^, a non-fluorescent version of GFP (**Fig. 2f** and **Supplementary Fig. 12**). The high affinity of ^GFP^LAMAs for GFP also allows to mislocalize GFP fusion proteins that are part of larger protein complexes. For example, transient transfection of mito-^GFP^LAMA_F98_ into a genome-edited cell line expressing NUP62-mEGFP (**Supplementary Fig. 13a**), a component of the nuclear pore complex, resulted in sequestering of NUP62-mEGFP from the nuclear envelope to the mitochondria in the absence of TMP (**Fig 2g** and **Supplementary Fig. 13b**). Upon addition of TMP, NUP62-mEGFP localized to the nuclear membrane.

GFP fusion proteins are omnipresent in the life sciences and our ^GFP^LAMAs offer a new way to probe the function of these proteins. To demonstrate the potential of ^GFP^LAMAs for mechanistic studies, we used mito-^GFP^LAMA_F98_ to control the function of a GFP fusion of Mad2L1, an important component of the mitotic checkpoint complex (**Fig 2h** and **Supplementary Fig. 14**). Knock-down of Mad2L1 reduces mitotic duration and increases the percentage of polylobed nuclei^19^. A HeLa Kyoto cell line in which endogenous Mad2L1 has been tagged with EGFP has been previously described and used to map the localizations of Mad2L1 during mitosis^20^. We stably expressed mito-^GFP^LAMA_F98_ in the Mad2L1-EGFP cell line, and observed how sequestering Mad2L1-EGFP to the mitochondria affected the outcome of cell division (**Supplementary Fig. 15** and **Supplementary Video 1-4**). In the absence of TMP, we observed an increase in the percentage of polylobed nuclei following mitotic events relative to cells not expressing mito-^GFP^LAMA_F98_ (85 ± 13 % *vs.* 10 ± 6 %; **Fig. 2i** and **Supplementary Fig. 16**). Addition of TMP to cells expressing mito-^GFP^LAMA_F98_ reduced the levels to those not expressing mito-^GFP^LAMA_F98_ (10 ± 11 %). Next, nocodazole, a small molecule which prevents attachment of microtubules to kinetochores, was added to activate the mitotic checkpoint complex. After treatment with nocodazole, cells in which Mad2L1-EGFP had been sequestered at the mitochondria were able to override mitotic arrest whereas treatment with nocodazole and TMP lead to mitotic arrest, as expected (**Fig. 2j** and **Supplementary Fig 17**). These data show that the function of Mad2L1-EGFP in the mitotic checkpoint complex can be controlled through its TMP-dependent interaction with mito-^GFP^LAMA_F98_.

In summary, LAMAs are a generally applicable chemogenetic tool to reversibly control the location and function of proteins, including the most commonly used class of fusion proteins (GFP). This tool opens up countless applications in research to study basic biological questions. As TMP is a clinically approved drug the approach might also be applicable *in vivo*. Furthermore, the design principle introduced here should be applicable for the generation of other switchable proteins.

## Acknowledgments

This work was supported by the Max Planck Society, the École Polytechnique Fédérale de Lausanne, the NCCR Chemical Biology. Research in the Kräusslich group was supported by the Deutsche Forschungsgemeinschaft (DFG, German Research Foundation) - Projektnummer 240245660 – SFB 1129 project 5 (H.-G.K). Research in the Ellenberg group was supported by the Paul G. Allen Frontiers Group through an Allen Distinguished Investigators Grant to J.E., the National Institutes of Health Common Fund 4D Nucleome Program (Grant U01 EB021223 / U01 DA047728 to J.E.) and the European Molecular Biology Laboratory (EMBL; S.O., M.K., J.E.). The authors thank Ilme Schlichting for X-ray data collection. Diffraction data were collected at the Swiss Light Source, beamline X10SA, of the Paul Scherrer Institute, Villigen, Switzerland. The authors would like to thank Luc Reymond, Johannes Broichhagen and Bettina Mathes for providing reagents. The authors would like to thank Manuel Eguren for valuable discussions.

## Additional Information

Requests for reagents and plasmids should be directed to Kai Johnsson. All requests for the Nup62-mEGFP cell line should be directed to Jan Ellenberg.

## Author Contributions

H.F. and K.J. designed the study. H.F generated, characterized and applied all LAMAs. M.T. solved the crystal structures of ^GFP^LAMAs. J.H. helped analyse the crystal structures. M.K. generated the NUP62-mEGFP cell line and S.O. performed the NUP62-mEGFP translocation experiments. B.K. helped with generation of stable cell lines with LAMAs. T.G.M. generated stable cells lines of ^p24^LAMA and characterized them. H.-G.K., J.E. and K.J. supervised the work. H.F and K.J. wrote the manuscript with input from all authors.

## ONLINE METHODS

### DNA plasmids and molecular cloning

Plasmids were generated using standard molecular biology techniques. All subcloned sequences were verified using Sanger sequencing, assisted by Geneious software (Biomatters). pCHIV is a non-infectious HIV-1 viral construct lacking LTRs and the *nef* gene^21^. pCHIV env(stop) was described earlier^22^ and contains a Klenow Polymerase fill in at the NdeI site to generate a frameshift within the *env* gene. mCherry was inserted between MA and CA of the gag polyprotein and was described earlier^23^. pWPI puro was obtained from Oliver Fackler^24^. psPAX2 was a gift from Didier Trono (Addgene plasmid #12260) and pCMV-VSV-G was a gift from Bob Weinberg (Addgene plasmid #8454).

### Chemical Reagents

Trimethoprim (TMP), pyrimethamine (Pyr) and methotrexate (MTX) were purchased from Sigma Aldrich. Stock solutions of 50 mM TMP in DMSO were used for both *in vitro* and in cell analysis. NADPH was purchased from PanReac Biochem, and stocks made to 10 mM fresh in aqueous buffer and used at the indicated concentrations. Nocodazol and reversine were purchased from Selleckchem. DMSO stock solutions were made fresh before use. Fluorescent dyes for live-cell imaging were purchased from available suppliers, or were synthesized as previously described^25,26^.

### Protein purification

Proteins were expressed using a pET51b(+) (Novagen) in *Escherichia coli* BL21 (DE3) pLysS, in the presence of 100 *µ*g mL^-1^ ampicillin in Luria-Bertani, shaking at 220 rpm. Cultures were grown at 37 °C until an OD_600_ of 0.8 was reached, and then induced with 1 mM isopropyl β-thiogalacopyranoside (ITPG). After overnight expression at 25 °C, cells were harvested and lysed by sonication. The lysates were cleared by centrifugation and purified by IMAC using Ni-NTA Resin (Thermo Fisher Scientific). For proteins used in TR-FRET assays, His-tag purification was followed by Strep-Tactin purifications (IBA Lifesciences), according to the manufacturer’s protocol.

### Fluorescence emission titrations

wtGFP emission assays were performed in 50 mM HEPES, 50 mM NaCl, 0.5 mg mL^-1^ BSA, 0.05% Triton-X 100, pH 7.3, in 384-well plates (Black, flat-bottom, Corning #3821). Dilution series of the protein switch in the presence of TMP (500 *µ*M, final concentration) or DMSO and/or NADPH (100 *µ*M, final concentration) were prepared and incubated at room temperature for 10 minutes. wtGFP (10 nM, final concentration) was diluted in 50 mM HEPES, 50 mM NaCl, 0.5 mg mL^-1^ BSA, 0.05% Triton-X 100, pH 7.3. Protein were then mixed in a 1:1 ratio in the plate, and the plate read in a Spark ® 20M microplate reader (Tecan). Excitation wavelength was 470 nm (5 nm bandwidth). Emission wavelength was 535 nm (5 nm bandwidth). The titration curves were fit with the full equation of single site binding, accounting for the effect of nonspecific binding^27^.

### TR-FRET assay

Constructs assayed by TR-FRET were expressed as SNAP-tag fusions and EGFP-fusions. SNAP-tag on the target proteins (4 *µ*M) was labeled with excess of SNAP-Lumi4-Tb (Cisbio) (6 *µ*M) in 50 mM HEPES, 50 mM NaCl, pH 7.3, at room temperature for 4 hours. Excess unlabeled dye was removed by centrifugal filter units (Amicon). Tb-labeled target protein was diluted in 50 mM HEPES, 50 mM NaCl, 0.5 mg mL^-1^ BSA, 0.05% Triton-X 100, pH 7.3, containing 100 *µ*M NADPH and placed into 384-well plates (Black, flat-bottom, Corning #3821). EGFP-fused proteins were diluted in 50 mM HEPES, 50 mM NaCl, 0.5 mg mL^-1^ BSA, 0.05% Triton-X 100 in the presence of TMP (500 *µ*M, final concentration) or DMSO and/or NADPH (100 *µ*M, final concentration). EGFP-fused proteins were mixed in a 1:1 ratio in the plate with the Tb-labeled target and incubated at least 15 min at room temperature. The plate was then read by a Spark ® 20M microplate reader (Tecan) in TR-FRET mode. The excitation wavelength was 320 nm (25 nm bandwidth). The emission wavelength for Tb was 480 nm (7.5 nm bandwidth), and the emission wavelength for EGFP was 520 nm (7.5 nm bandwidth) using a 510 dichroic mirror. Integration time was 400 *µ*s and lag time 120 *µ*s. The titration curves were fit the full equation of single site binding, accounting for the effect of nonspecific binding.

### Kinetics of in vitro dissociation

wtGFP was mixed with an excess of LAMA (1:3 ratio) on ice for 10 min and passed over size-exclusion chromatography. The heterodimeric fraction was collected and diluted into 50 mM HEPES, 50 mM NaCl, 0.5 mg mL^-1^ BSA, 0.05% Triton-X 100, pH 7.3, in 96-well black flat-bottomed plates to a final concentration of 200 nM, in the presence of 100 *µ*M NADPH. TMP, Pyr or MTX were diluted in DMSO. 1 *µ*L of relevant drug solutions was added to the 96-well plate and fluorescence emission recorded over time on a Spark ® 20M microplate reader (Tecan). Excitation wavelength was 470 nm (5 nm bandwidth). Emission wavelength was 535 nm (5 nm bandwidth). For reversible association eDHFR was diluted in 50 mM HEPES, 50 mM NaCl, 0.5 mg mL^-1^ BSA, 0.05% Triton-X 100 and 1 *µ*L added to the reaction mix in the 96-well plate. The curves where fit with one-phase dissociation models to estimate the half time at these concentrations.

### Protein crystallization

For X-ray crystallography, the LAMAs were sub-cloned into a vector carrying an N-terminal Hisx10-tag, followed by a tobacco etch virus (TEV) protease cleavage tag sequence. The production and IMAC purification of the TEV protease was performed as previously described^28^. The His-Tag was removed from the LAMAs by TEV protease cleavage at 30 °C overnight, at a ratio of 1:20 (TEV protease: LAMA). The digested protein was purified using a reverse IMAC purification method by NTA resin, collecting the flow-through. The protein was passed over a size-exclusion column and concentrated using centrifugal filter units (Amicon). The protein was flash-frozen and stored at ℒ70 °C. Purified protein in 25 mM HEPES, 25 mM NaCl, pH 7.3 was premixed with NADPH (10 eq) and TMP (10 eq) as solid powders in 300 *µ*L volume. The solution was left on ice for 10 minutes before centrifugation (20 000*g*, 10 min, 4 °C) to remove any precipitation.

Crystallization was performed at 20°C using the vapor-diffusion method. Crystals of ^GFP^LAMA_F98_:NADFPH:TMP complex with a rod morphology were grown by mixing equal volumes of protein solution at 25 mg/ml in 25 mM HEPES, 25 mM sodium chloride pH 7.3 and a reservoir solution containing 0.1 M MES pH 6.0, 30% (v/v) PEG 600, 5% (w/v) PEG 1000 and 10% (v/v) glycerol. The crystals were briefly washed in cryoprotectant solution consisting of the reservoir solution with sucrose and glucose added to a final concentration of 10% (w/v) each, prior to flash-cooling in liquid nitrogen. ^GFP^LAMA_G97_ :NADPH:TMP complex crystals were obtained by mixing equal volumes of protein solution at 15 mg/ml in 25 mM HEPES, 25 mM sodium chloride pH 7.3 and precipitant solution containing 0.1 M MES pH 6.0, 20% (w/v) PEG 6000 and 1.0 M lithium chloride. Thin plate-shaped crystals grew in clusters; single plates could be isolated and were briefly washed in cryoprotectant solution consisting of the reservoir solution supplemented with 20% (v/v) glycerol before flash-cooling in liquid nitrogen.

### X-ray diffraction data collection and structure determination

Single crystal X-ray diffraction data were collected at 100 K on the X10SA beamline at the SLS (PSI, Villigen, Switzerland). All data were processed with XDS^29^. The structures were determined by molecular replacement (MR) using Phaser^30^ and individual protein coordinates from PDB entries 5UII and 5H8D as a search models for DHFR and nanobody, respectively. The final models were optimized in iterative cycles of manual rebuilding using Coot^31^ and refinement using Refmac5^32^ and phenix.refine^33^. Data collection and refinement statistics are summarized in Supplementary Table 1, model quality was validated with MolProbity^34^ as implemented in PHENIX. The omit maps for ligands were generated using the composite omit map tool in PHENIX^33^.

Atomic coordinates and structure factors have been deposited in the Protein Data Bank under accession codes: 6RUL (^GFP^LAMA_F98_), 6RUM (^GFP^LAMA_G97_). Analysis was performed using MacPyMOL^35^ and Coot^31^.

### Mammalian cell culture maintenance

Eukaryotic cells were obtained from American Type Culture Collection (ATCC, Manassas, Virginia), The Leibniz Institute DSMZ-German Collection of Microorganisms and Cell Cultures (DSMZ, Germany), or from collaborators as indicated. No cell lines on the ICLAC list of commonly misidentified cells were used in this work. All cells were cultured in DMEM GlutaMax (Thermo Fisher Scientific) medium supplemented with 10% FBS, and penicillin and streptomycin as indicated, at 37 °C in a humidified incubator with 5% CO_2_. All cells were mycoplasma-free.

### Generation of ^p24^LAMA_s98_ cell lines

Lentiviral particles were generated by co-transfection of transfer plasmid pWPI EGFP-^p24^LAMA_S98_ IRES puro, packaging construct psPAX2, fusion protein expression plasmid pCMV-VSVG and pAdvantage (Promega) in a ratio of 1.5 : 1 : 0.5 : 0.2, into HEK293T (ATCC) cells using PEI (1:3 ratio of *µ*g DNA : *µ*l 1 mg/ml PEI). The medium was changed after 6 hours and production of lentiviral particles was allowed to proceed for 48 h. The supernatant was filtered through 0.45 *µ*m MCE filters and was directly added to HeLa TZM-bl cells (NIH AIDS Repository). After 2 days the cells were expanded and 1 *µ*g/ml puromycin was added to select for stably transduced cells.

### Genome editing

Nup62 in HeLa Kyoto cells was endogenously tagged with mEGFP at the C-terminus by CRISPR-Cas9 nickases and its homozygous integration was validated as described previously^36,37^. The gRNA sequences for the genome editing are as follows: 5’TCGCTCAGTCAAAGGTGATC3’ and 5’CTGGGGCCCGCAGGTCCCTA3’.

Previously described genome edited HeLa Kyoto Mad2L1-EGFP^20^ were used in the generation of stable cell lines expressing LAMAs. HeLa Kyoto Mad2L1-EGFP cells were seeded one day before transfection with Lipofectamin3000 (Thermo Fisher Scientific), in the presence of TMP (10-50 *µ*M), following the manufacturer’s instructions. Cells were seeded monoclonal densities and selected using 500-800 *µ*g mL^-1^ geneticin (Thermo Fisher Scientific), and TMP (10-50 *µ*M) for 2 weeks. Cells were then labeled with BG-SiR (500 nM) overnight, before sorting on a FACSMelody (BD biosciences) for LAMA expressing cells.

### Live cell imaging of ^p24^LAMA_S98_ in HIV expressing cells

HeLa TZM-bl cells were seeded the day before transfection in complete medium with 100 U/ml penicillin 100 *µ*g/ml streptomycin and incubated at 37 °C and 5 % CO_2_. pCHIV env(stop) was transfected in a 1:1 ratio with pCHIV env(stop) gag-mCherry using Turbofect (1:2 ratio). The cells were incubated for 24 hours, the medium was changed to imaging medium (FluoroBrite DMEM (Thermo Fisher Scientific), 10 % FBS, 4 mM GlutaMAX (Gibco Life Technologies), 2 mM sodium pyruvate (Gibco Life Technologies), 20 mM HEPES pH 7.4, 100 U/ml Penicillin 100 *µ*g/ml Streptomycin (PAN-Biotech, Germany)) and transferred to a Nikon Eclipse Ti2 (Nikon, Japan) inverted microscope equipped with an Andor confocal spinning disc unit (Yokogawa CSU-W1 Spinning Disk Unit, Andor, Oxford Instruments, United Kingdom). Cells were imaged at 37 °C and 5 % CO_2_ using a 100 × oil-immersion objective (Nikon CFI Apochromat TIRF 100X Oil NA 1.49) and a dual EMCCD camera setup (ANDOR iXon DU-888), simultaneously recording the EGFP (488/500-550 nm) and the mCherry channel (568/575-625 nm) with a pixel size of 0.13 *µ*m. 3D stacks (0.5 *µ*m, z-spacing) were recorded with a time interval of 3 minutes for 60-120 minutes at up to 32 randomly chosen positions using the Nikon Imaging Software Elements 5.02. After 4-12 frames 50 *µ*l TMP containing imaging medium was added to a final concentration of 10 *µ*M and imaging was continued. The movies were filtered in Fiji/ImageJ with a mean filter (kernel size: 0.25 × 0.25 *µ*m) to reduce noise and the mean intensity of a region of interest inside the nucleus was measured using the Multi Measure function of the ROI Manager. Camera background was subtracted and intensities where normalized to the highest intensity within the ROI during the timeseries to correct for different expression levels. Different experiments were temporally aligned to the time of TMP addition and data was pooled from 3 independent experiments.

### Translocation of fluorescent proteins

HeLa Kyoto cells^36^ were seeded 24 hours before transfection with Lipofectamin2000 (Thermo Fisher Scientific) according to the manufacturer’s protocol. After 24 hours, complete medium without phenol red was added to the cells. Cells were labeled with BG-SiR (500 nM) overnight, in the presence or absence of 10 *µ*M TMP or DMSO, before being imaged by confocal microscopy using a Leica DMi8 microscope (Leica Microsystems, Germany) equipped with a Leica TCS SP8 X scanhead; a SuperK white light laser, and an HC PL APO 40×/1.10 W motCORR CS2 objective, at 37 °C with 5 % CO_2_, achieved by a temperature controllable incubator (Life Imaging Services). Image acquisition was performed with speed of 400 Hz, pixel dwell time 600 ns, pixel size 0.06 *µ*m, with z-stacks of 1 *µ*m over 10 *µ*m. A white-light laser was used for excitation, collecting with Leica HyD detectors: EGFP (488/505-550 nm), YFP (514/525-573 nm), ShadowG-mScarlet (561/583-625 nm), SiR (633/650-750 nm).

### Perfusion of TMP over live cells

U-2 OS (DSMZ) cells were transfected with Lipofectamin2000 (Thermo Fisher Scientific) according to the manufacturer’s protocol. After 24 hours, cells were seeded on Ibidi 0.6 Luer I cell culture treated perfusion chambers. After the cells were adherent, BG-SiR (500 nM) was added to the perfusion chamber and labeled overnight at 37 °C. The perfusion chamber was then attached to a custom-built gravity-perfusion system and mounted on a Leica DMi8 microscope (Leica Microsystems, Germany) equipped with a Leica TCS SP8 X scanhead; a SuperK white light laser, and an HC PL APO 40×/1.10 W motCORR CS2 objective, at 37 °C. TMP (0.5 *µ*M-20 *µ*M) in complete DMEM GlutMax medium with phenol red was perfused over the cells, and images acquired at a scanning speed of 400 Hz, pixel dwell time 1.2 *µ*s, with a pinhole at 1 airy unit, with z-stacks of 1 *µ*m over 10-20 *µ*m, with image acquisition every 30-60 s. A white-light laser was used for excitation, collecting with Leica HyD detectors: EGFP (488/505-550 nm), SiR (633/650-750 nm). Image analysis was performed in Fiji/ImageJ using the Time Series Analyzer (3.0), selecting a ROI of interest in the nucleus, and measuring the fluorescent intensity over time, with background subtraction of a region outside of the cells. All values were normalised to the intensity in the nucleus after the highest concentration of TMP perfused for each cell. Different experiments were temporally aligned to the time of TMP addition and data was pooled from 3 independent experiments.

### Live-cell imaging of Nup62-mEGFP

Live-cell imaging of HeLa Nup62-mEGFP cells was performed at 37 °C in CO_2-_ independent medium without phenol red (Invitrogen, Carlsbad, CA) containing 20% FBS, 2 mM l-glutamine, and 100 *µ*g/ml penicillin and streptomycin, with either 10 *µ*M of TMP or DMSO. Cells were incubated with 10 *µ*M BG-TMR for 30 min and the BG-TMR was washed away before imaging. Cells were then observed by confocal microscopy (LSM780; Carl Zeiss, Oberkochen, Germany) using a 63 × 1.4 NA Plan-Apochromat objective (Carl Zeiss), recording the mEGFP (488/491-552 nm) and TMR (561/580-660 nm) channels with a xy resolution of 0.13 *µ*m and the section thickness of 1.2 *µ*m. Fluorescence images were filtered with a median filter (kernel size: 0.25 × 0.25 *µ*m) for presentation purposes.

### Automated microscopy and analysis

For continuous live-cell imaging of HeLa Kyoto Mad2L1-EGFP cells stably expressing LAMAs, cells were seeded in 96-well plates (Eppendorf), in the presence of TMP (50 *µ*M). After cells were adherent, cells were labeled with BG-SiR (100 nM) overnight. The cells were then labeled with Hoechst 33342 (1 *µ*g/ mL) in complete medium without phenol red in the presence or absence of TMP (50 *µ*M) for 15 min, and washed 3 times with complete medium without phenol red in the presence or absence of TMP (50 *µ*M). Cells were imaged in the presence of BG-SiR (100 nM), in the presence or absence of TMP (50 *µ*M), and in the presence of additional mitotic drugs as indicated, nocodazole (330 nM) or reversine (5 *µ*M). Automatic microscopy was performed with Leica HCS A Matrix Screener software on a Leica DMi8 microscope (Leica Microsystems, Germany) equipped with a Leica TCS SP8 X scanhead; a SuperK white light laser, and an HC PL APO 40×/1.10 W motCORR CS2 objective, at 37 °C, 5% CO_2_, achieved by a temperature controllable incubator (Life Imaging Services). A white-light laser or 405 nm diode was used to excite the fluorophores, collecting with Leica HyD detectors: Hoechst (405/425-475 nm), SiR (633/650-750 nm). Image acquisition was performed with speed of 400 Hz, pixel dwell time 1.2 *µ*s, pixel size 0.57 *µ*m, with z-stacks of 1 *µ*m over 10 *µ*m. Image analysis was performed in Fiji/ImageJ, with manual annotations of LAMA expressing cells followed from prometaphase to mitotic exit.

