## Supplementary Information for "Chemogenetic Control of Nanobodies"

**Table of contents**

**Supplementary Fig. 1.** Generation of LAMAs from the enhancer nanobody 3

**Supplementary Fig. 2.** Complex formation between $^{GFP}$LAMAs and wtGFP 4

**Supplementary Fig. 3.** ITC measurements of the interaction between LAMAs and wtGFP 5

**Supplementary Fig. 4.** Kinetics of dissociation and association between LAMAs and wtGFP 6

**Supplementary Fig. 5.** Structural analysis of $^{GFP}$LAMAs 7

**Supplementary Fig. 6.** Titrations of $^{GFP}$LAMAs with GGS-linkers 8

**Supplementary Fig. 7.** Insertion of cpDHFR into other nanobodies 9

**Supplementary Fig. 8.** Generation of LAMAs from the p24 binding nanobody, cb9 10

**Supplementary Fig. 9.** Characterization of $^{p24}$LAMA$_{S98}$ in live cells 11

**Supplementary Fig. 10.** Live cell imaging of localized $^{GFP}$LAMAs with EGFP 12

**Supplementary Fig. 11.** Perfusion experiments in U-2 OS cells with mito-$^{GFP}$LAMA$_{F98}$ 13

**Supplementary Fig. 12.** Live cell imaging of GFP-LAMAs extended to YFP and ShadowG-mScarlet 14

**Supplementary Fig. 13.** Use of $^{GFP}$LAMAs to mislocalize NUP62-mEGFP 15

**Supplementary Fig. 14.** Mislocalization of Mad2L1-EGFP using $^{GFP}$LAMAs 16

**Supplementary Fig. 15.** Cellular trajectories of live cell imaging of Mad2L1-GFP cells stably expressing $^{GFP}$mitoLAMA$_{F98}$ 17

**Supplementary Fig. 16.** Nuclear morphology of Mad2L1-EGFP after mislocalization 18

**Supplementary Fig. 17.** Duration of mitosis in Mad2L1-EGFP cells expressing mito-$^{GFP}$LAMA$_{F98}$ 19

**Supplementary Videos 1-4.** Live cell imaging of Mad2L1-GFP expressing a mito-LAMA 20

**Supplementary Table 1.** X-ray data collection and refinement statistics 21

**Amino acid sequences of LAMAs used** 22

**Additional notes, equations and probe sequences** 23

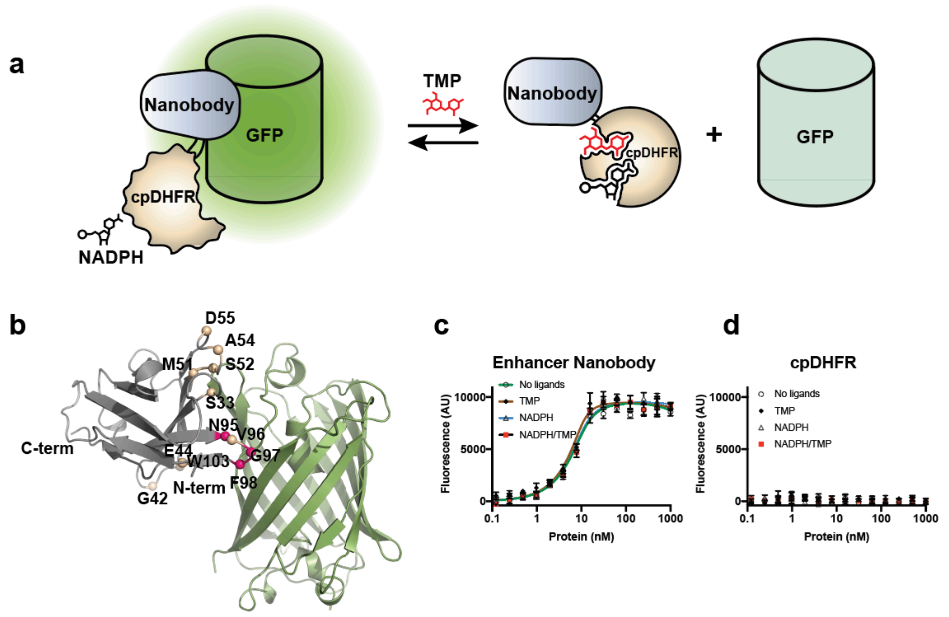

**e**

| Insertion position | $K_d$ No TMP (nM) | $K_d$ TMP (nM) | Region of nanobody | Switch |
|---|---|---|---|---|
| **None** | Below 1 | Below 1 | - | No |
| **S33** | 20 ± 5 | 39 ± 11 | CDR1 | No |
| **G42** | 45 ± 6 | 49 ± 7 | FR2 | No |
| **E44** | 109 ± 14 | 147 ± 17 | FR2 | No |
| **$M51_{GGS}$** | 1.6 ± 0.3 | 50 ± 5 | CDR2 | Slight |
| **S52** | 40 ± 3 | 170 ± 16 | CDR2 | Slight |
| **A54** | 1.7 ± 0.4 | 2.3 ± 0.5 | CDR2 | No |
| **D55** | Below 1 | Below 1 | CDR2 | No |
| **N95** | 3.3 ± 0.5 | No fit | CDR3 | Yes |
| **V96** | 99 ± 14 | No fit | CDR3 | Yes |
| **G97** | 6.0± 0.5 | No fit | CDR3 | Yes |
| **F98** | 79 ± 7.3 | No fit | CDR3 | Yes |
| **W103** | Approx. 500 | Approx. 500 | FR4 | No |

**Supplementary Fig. 1.** Generation of LAMAs from the enhancer nanobody. (**a**) Schematic illustration of the $^{GFP}$LAMAs. The enhancer nanobody enhances the fluorescent emission of wtGFP when bound to it. (**b**) Insertion sites of cpDHFR mapped onto the X-ray structure of the enhancer nanobody (grey) bound to GFP (green) (PDBID = 3K1K) as spheres. Insertion positions with large potential for switching are highlighted in magenta. (**c,d**) Titration between recombinant enhancer nanobody (**c**) or cpDHFR (**d**) and wtGFP, as quantified by fluorescent emission increase of wtGFP. Mean ± s.d. representative from 3 independent experiments. NADPH = 100 µM, TMP = 500 µM. (**e**) Table of affinities from the titrations between the enhancer nanobody with cpDHFR insertions and wtGFP. Data were fit the full equation of single site binding, accounting for the effect of nonspecific binding, with s.e.m. of the fit. Insertion position is represented as the last residue prior to cpDHFR insertion.

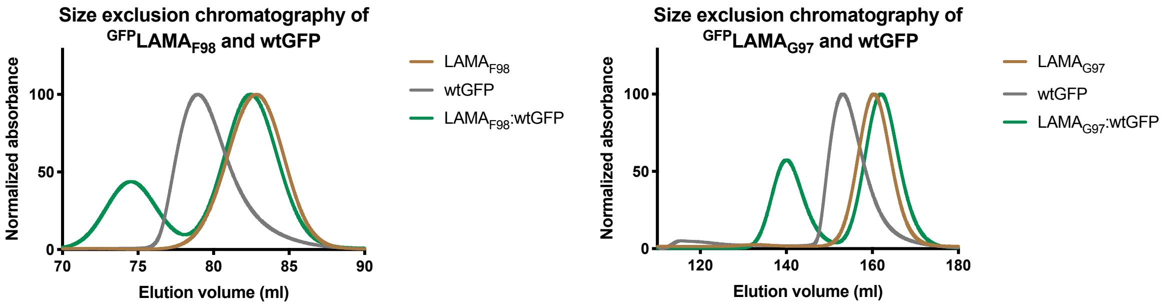

**Supplementary Fig. 2.** Complex formation between $^{GFP}$LAMAs and wtGFP. Size exclusion chromatography was performed with LAMAs and wtGFP. Absorbances are normalized (%) to the maximum intensity values for each run. LAMAs and wtGFP were each injected separately on the column for calibration. An excess of $^{GFP}LAMA_{F98}$ or $^{GFP}LAMA_{G97}$ (2-5 fold) was premixed with wtGFP and incubated on ice for 10 minutes, before being separated on the same column. For $^{GFP}LAMA_{F98}$ an S200-16-600 (GE-Healthcare) was used. For $^{GFP}LAMA_{G97}$ an S75-26-600 (GE-Healthcare) was used.

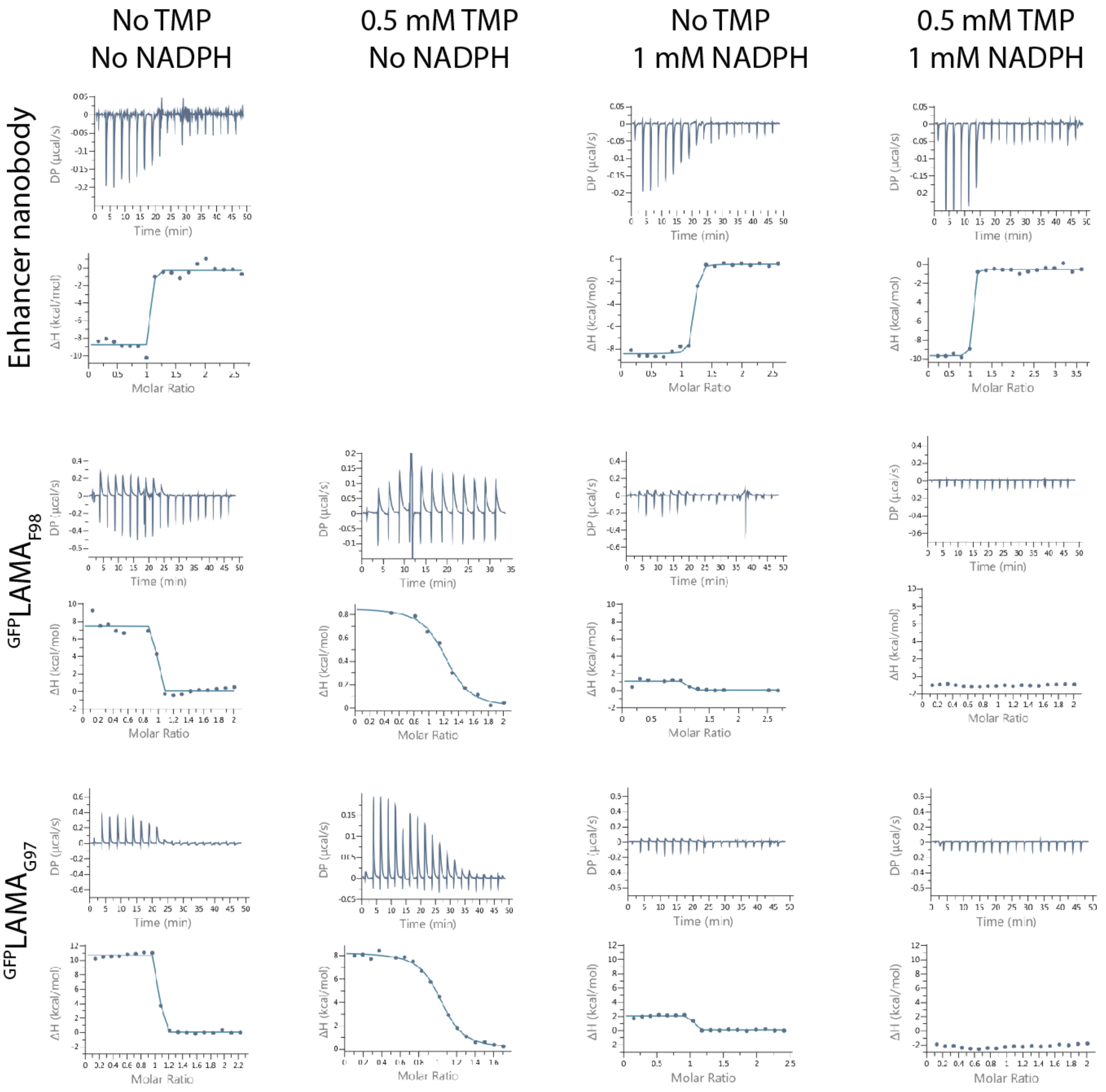

| LAMA | $K_d$ No Ligands (nM) | $K_d$ TMP (nM) | $K_d$ NADPH (nM) | $K_d$ TMP/NADPH (nM) |
|---|---|---|---|---|
| **Nanobody only** | Below 1 | Below 1 | Below 1 | Below 1 |
| **F98** | 2.3 ± 0.2 | 712 ± 57 | 79 ± 7.3 | No fit |
| **G97** | 1.2 ± 0.3 | 289 ± 17 | 6.0 ± 0.5 | No fit |
| **V96** | 34 ± 6 | 1300 ± 300 | 99 ± 14 | No fit |
| **N95** | 3.2 ± 0.5 | 13 ± 1 | 3.3 ± 0.5 | No fit |

**Supplementary Fig. 3.** ITC measurements of the interaction between LAMAs and wtGFP. Proteins were dialysed against buffers containing NADPH (1 mM) and/or TMP (0.5 mM), before measurements. The data was fit using the Malvern PEAQ-ITC analysis software, fixing the binding as 1:1 and calculating a $K_d$ and an error in the fit. Representative traces of 3 independent experiments.

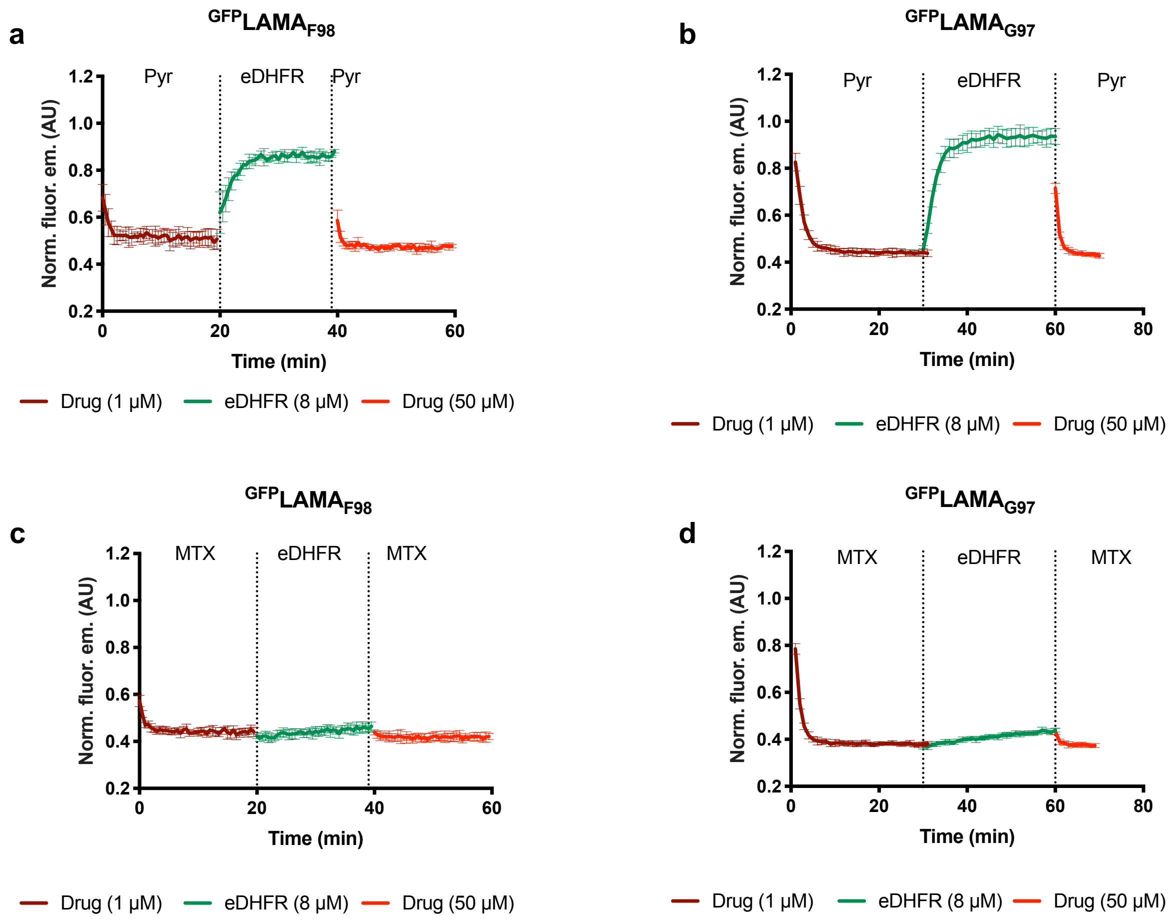

**Supplementary Fig. 4.** Kinetics of dissociation and association between LAMAs and wtGFP. Addition of different dihydrofolate drugs was performed in the presence of NADPH (100 μM). Dissociation was followed by the decrease in fluorescent intensity of wtGFP on the addition of pyrimethamine (Pyr) (**a**,**b**), and methotrexate (MTX) (**c**,**d**). Mean ± s.d., representative of 2 independent experiments.

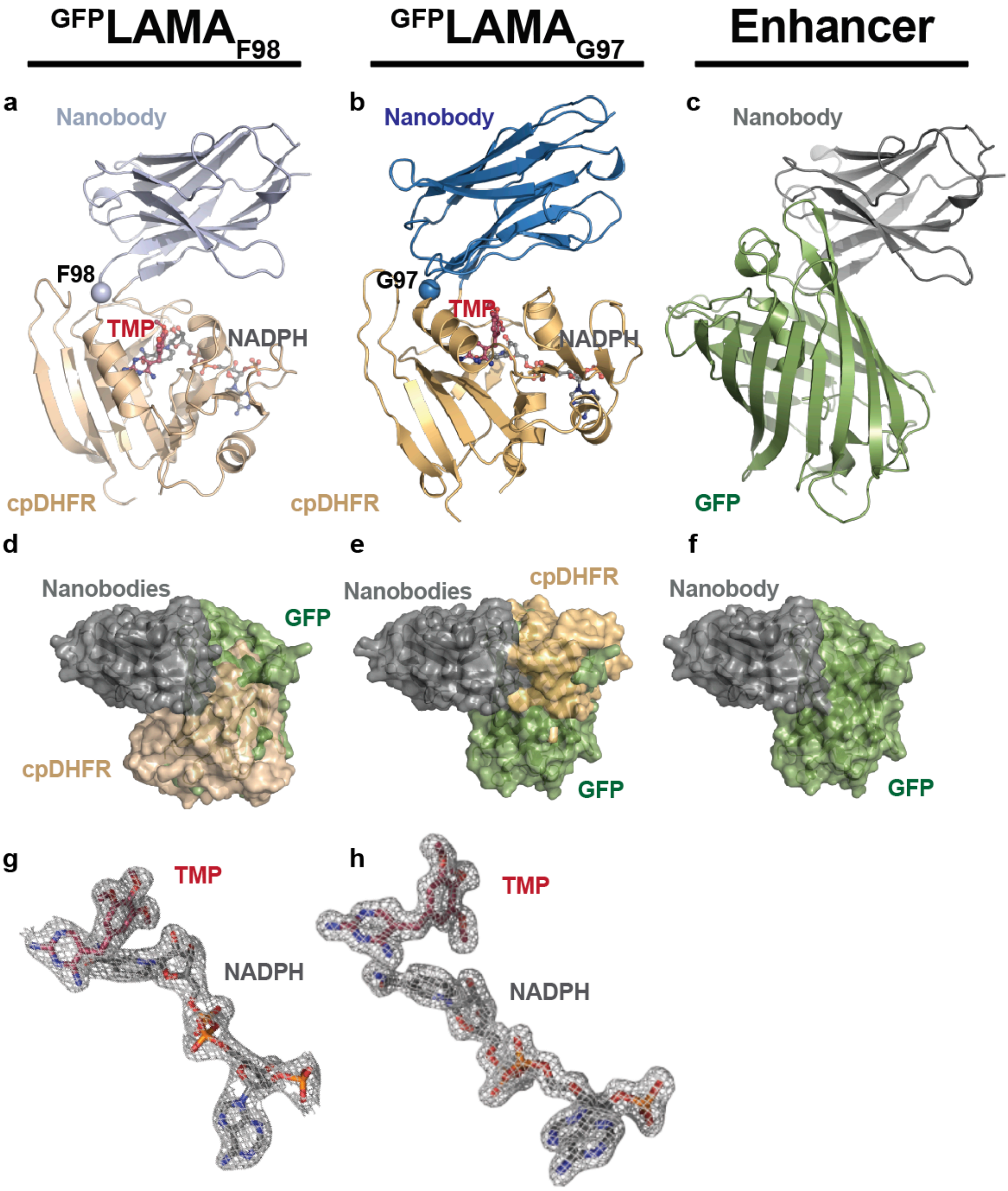

**Supplementary Fig. 5.** Structural analysis of $^{GFP}$LAMAs. (**a-c**) Comparison between the X-ray structures of $^{GFP}$LAMA$_{F98}$ (PDBID = XXX) (**a**), $^{GFP}$LAMA$_{G97}$ (PDBID = XXX) (**b**), both in the presence of TMP (red), NADPH (grey), and enhancer nanobody bound to GFP (PDBID = 3K1K) (**c**). Structures are represented as cartoons, ligands as stickballs. The insertion positions of cpDHFR are highlighted on the LAMA structures. (**d-f**) Overlay of the nanobody domain of LAMA structures on the enhancer nanobody bound to GFP with $^{GFP}$LAMA$_{F98}$ (**d**), and $^{GFP}$LAMA$_{G97}$ (**e**). The enhancer nanobody bound to GFP (**f**) is shown for comparison. (**g,h**) The 2mFobs-DFcalc omit electron density maps showing the placement of TMP and NADPH in cpDHFR of $^{GFP}$LAMA$_{F98}$ (**g**) and $^{GFP}$LAMA$_{G97}$ (**h**) contoured at 1.0 $\sigma$.

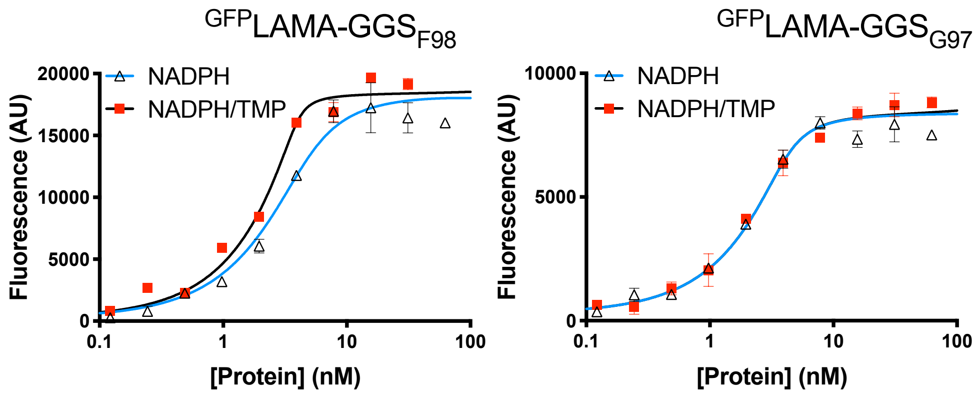

**Supplementary Fig. 6.** Titrations of $^{GFP}$LAMAs with GGS-linkers. Fluorescent emission from titration between cpDHFR inserted with GGS-linkers into the nanobody structure and wtGFP. Mean ± s.d, fit with the full equation of single site binding, accounting for the effect of nonspecific binding. Representative experiment from 2 trials.

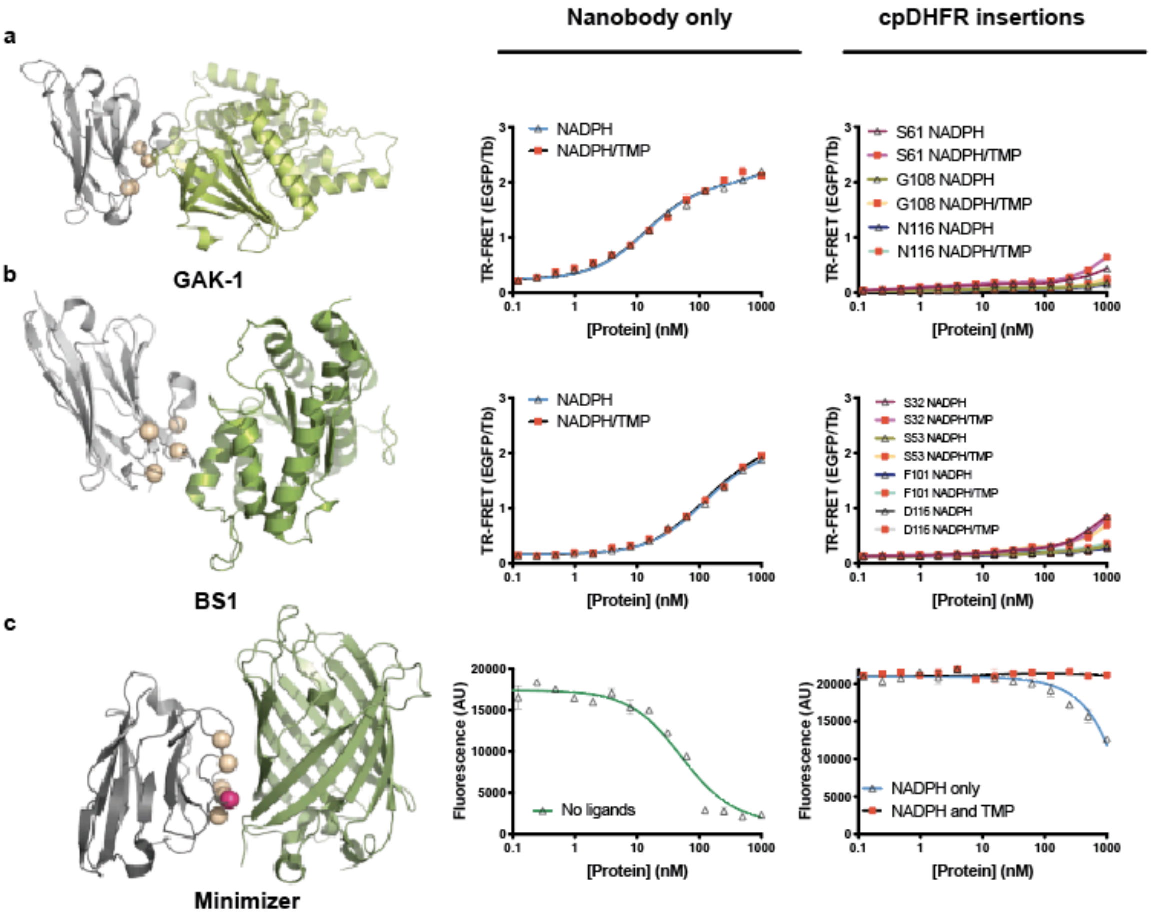

**Supplementary Fig. 7.** Insertion of cpDHFR into other nanobodies. Left panel: X-ray structures of nanobodies (grey) and targets (green) in complex. Insertion positions of cpDHFR tried are mapped as beige spheres, successful positions as magenta speheres. (**a**) LAMA strategy for anti-GAK nanobody (PDBID = 4C57) and (**b**) BS1 nanobody (PDBID = 4TVS). TR-FRET was used as a readout for affinity between the EGFP-fused nanobody and terbium-cryptate labelled target via SNAP-tag. (**c**) LAMA strategy for the minimizer nanobody (PDBID = 3G9A). Only data for ligand-dependent G100g insertion is plotted for clarity. Interaction is measured as a decrease in EGFP fluorescent intensity. Titrations are mean ± s.d. from duplicates.

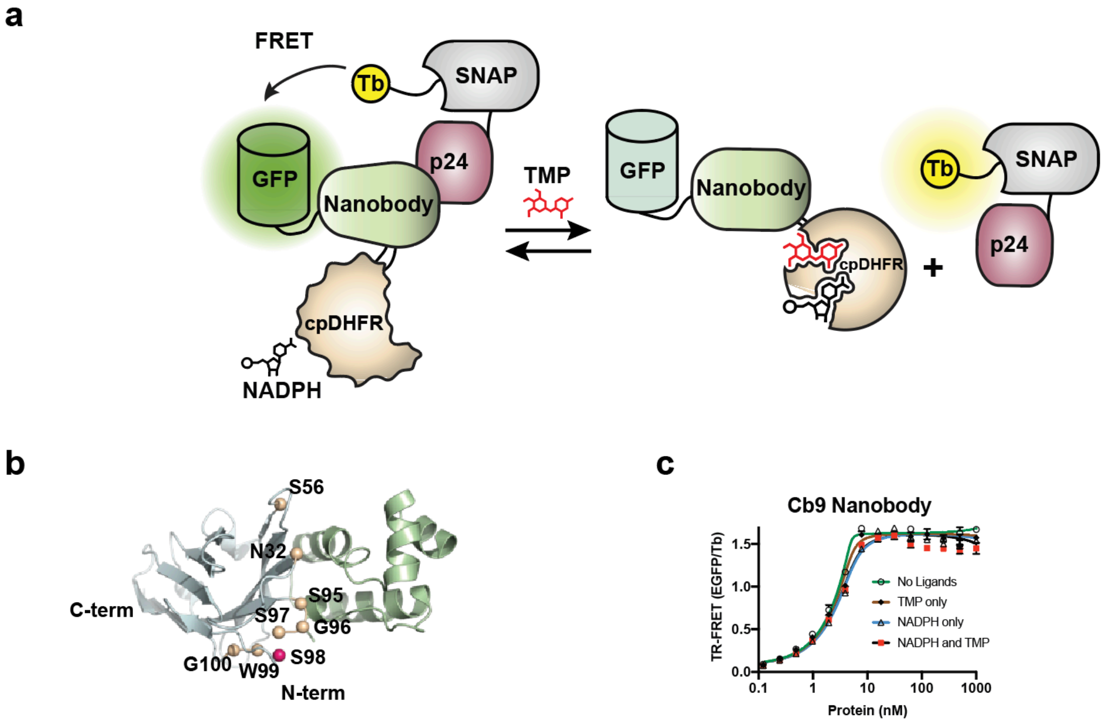

**d**

| Insertion position | NADPH only (nM) | $K_d$ NADPH and TMP (nM) | Region of nanobody | Switch |
|---|---|---|---|---|
| **None** | 1.1 ± 0.3 | 1.6 ± 0.3 | - | No |
| **N32** | Below 1 | Below 1 | CDR1 | No |
| **S56** | 1.4 ± 0.1 | Below 1 | CDR2 | No |
| **S95** | 3.5 ± 0.4 | 70 ± 18 | CDR3 | Yes |
| **G96** | 330 ± 10 | 2220 ± 50 | CDR3 | Yes |
| **S97** | 63 ± 2 | 1100 ± 100 | CDR3 | Yes |
| **S98** | 7 ± 1 | 524 ± 34 | CDR3 | Yes |
| **W99** | 22 ± 1 | 50 ± 3 | CDR3 | Limited |
| **G100** | 14 ± 1 | 18 ± 1 | CDR3 | No |

**Supplementary Fig. 8.** Generation of LAMAs from the p24 binding nanobody, cb9. (**a**) Schematic illustration of the TR-FRET assay to monitor binding between the cb9 nanobody and the p24 target protein. The $^{p24}$LAMA is expressed as an EGFP fusion (FRET acceptor) and the p24 target expressed as a SNAP-tag fusion, labelled with Tb-cryptate (FRET donor). (**b**) crystal structure of cb9 (blue) with the C-terminal region of p24 (green) represented as cartoon, with insertion sites of cpDHFR represented as spheres (PDBID = 2XV6). The insertion positions with a large potential for switching is highlighted in magenta. (**c**) Titration between recombinant cb9 nanobody and p24, measured by TR-FRET. Mean ± s.d., representative from 3 independent experiments. NADPH = 100 µM, TMP = 500 µM. (**d**) Table of affinities between the cb9 nanobody and p24. Data were fit with the full equation of single site binding, accounting for the effect of nonspecific binding, with s.e.m. of the fit. Insertion position is represented as the last residue prior to cpDHFR insertion.

**a**

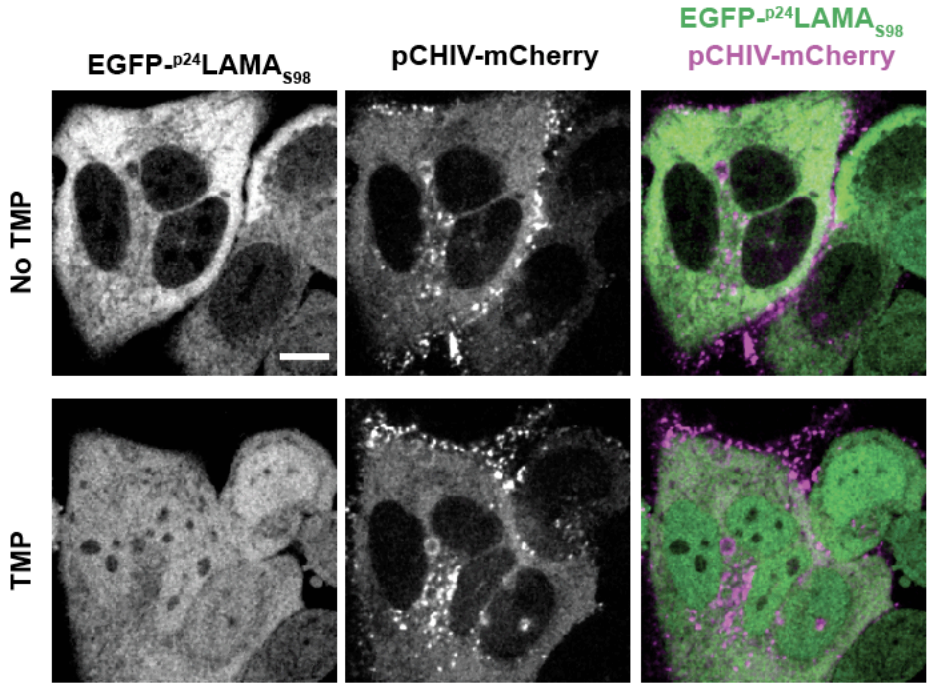

**b**

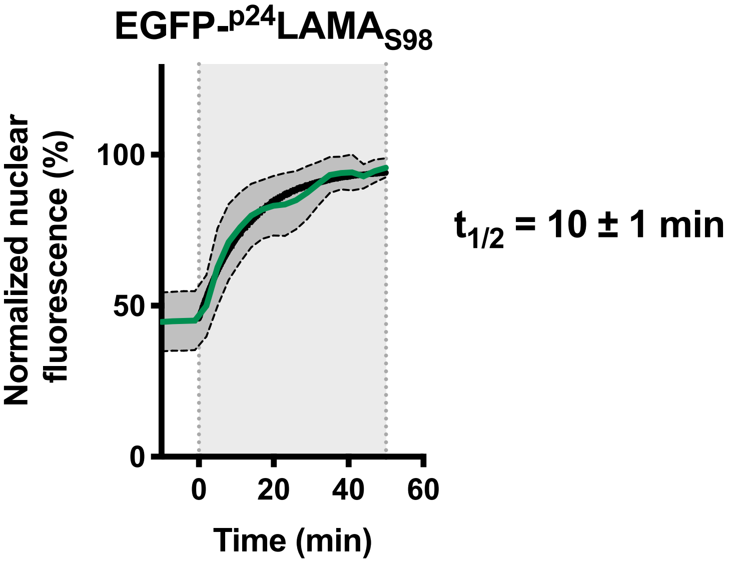

**Supplementary Fig. 9.** Characterization of $^{p24}$LAMA$_{S98}$ in live cells. (**a**) Full panel of micrographs from before and after addition of TMP (10 µM) to HeLa TZM-bl cells transfected with pCHIV-mCherry. EGFP-$^{p24}$LAMA$_{S98}$ , pCHIV-mCherry and merge are shown. Viral assembly sites of HIV are visualized as bright puncta in the mCherry channel. Scale bar, 10 µm. (**b**) Kinetics of nuclear EGFP-fluorescence after addition of TMP (10 µM, grey background), TMP was added at time point 0, mean ± s.d. are shown as green line and dashed bar with dark grey fill, from N = 46 cells from 3 independent experiments. Data was fit with a one-phase association curve, shown in black. $t_{1/2}$ is calculated from the 46 individual traces and stated as the mean ± s.e.m..

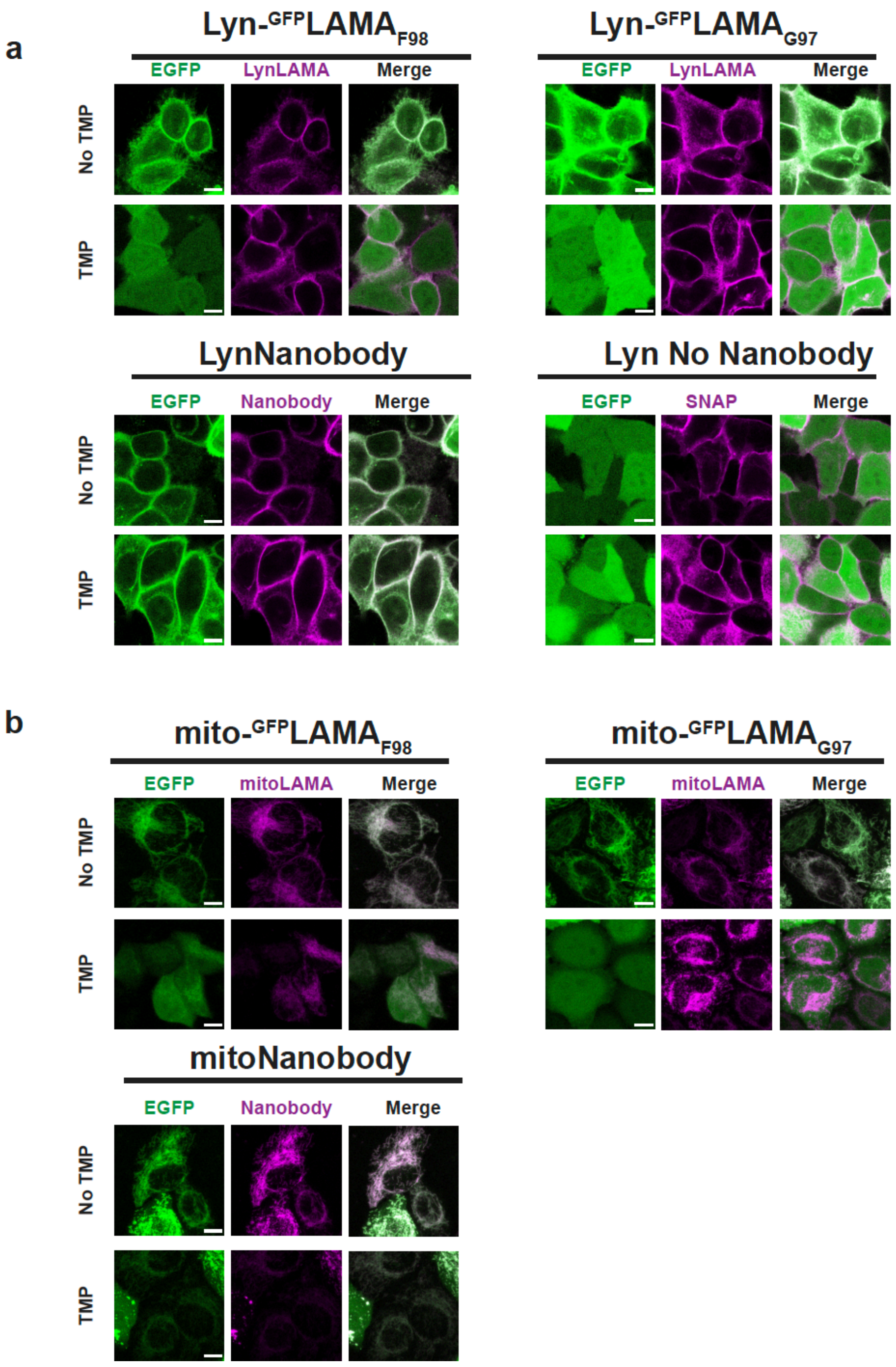

**Supplementary Fig. 10.** Live-cell imaging of localized $^{GFP}$LAMAs with EGFP. (**a**) LAMAs anchored to the inner plasma membrane through a Lyn-sequence. EGFP was coexpressed with no cellular localization. (**b**) LAMAs anchored to the outer mitochondrial membrane using a Ntom20 sequence. Lyn-and mitoNanobody indicates the enhancer nanobody with no cpDHFR insertion (positive control). No Nanobody indicates SNAP-tag anchored with no nanobody or LAMA (negative control). SNAP-tag was imaged after labelling with BG-SiR (500 nM). TMP = 10 µM. Scale bar, 10 µm.

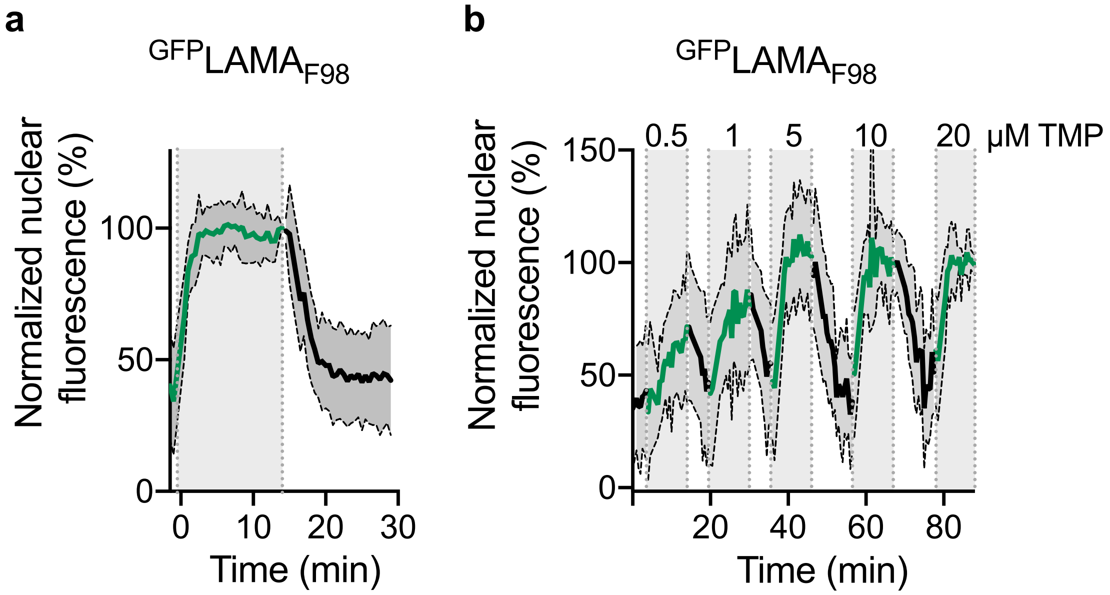

**Supplementary Fig. 11.** Perfusion experiments in U-2 OS cells with mito-$^{GFP}$LAMA$_{F98}$. The fluorescence of EGFP in the nucleus was quantified. (**a**) TMP (10 µM) perfused over cells (N = 24) at time point zero. Media was used to washout TMP, as indicated by the end of the grey zone. (**b**) Titration of increasing concentration of TMP perfused over cells (N = 21), with washout of media in between each perfusion. Mean ± s.d. indicated as a solid line and grey dashed area. Fluorescence emission was normalized for each cell's maximal fluorescence value after perfusion of TMP.

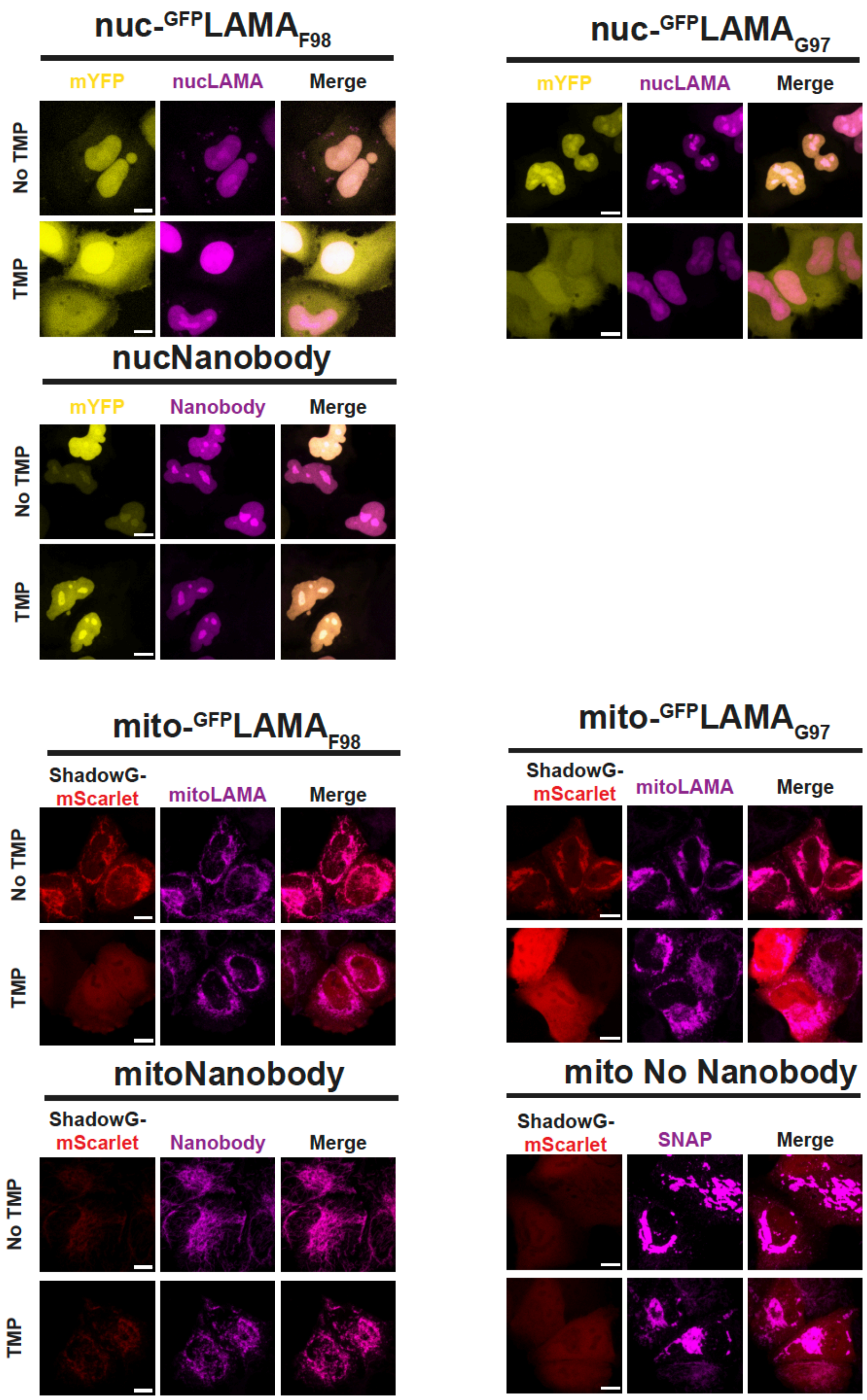

**Supplementary Fig. 12.** Live cell imaging of GFP-LAMAs extended to YFP and ShadowG-mScarlet. $^{GFP}$LAMAs with nuclear and mitochondrial localisation were coexpressed with YFP or mScarlet-ShadowG. mScarlet was fused to ShadowG for a marker in imaging. nuc-and mitoNanobody indicates the enhancer nanobody with no cpDHFR insertion (positive control). No Nanobody indicates SNAP-tag anchored with no nanobody or LAMA (negative control). SNAP-tag was imaged by labelling with BG-SiR (500 nM). TMP = 10 µM. Scale bar, 10 µm.

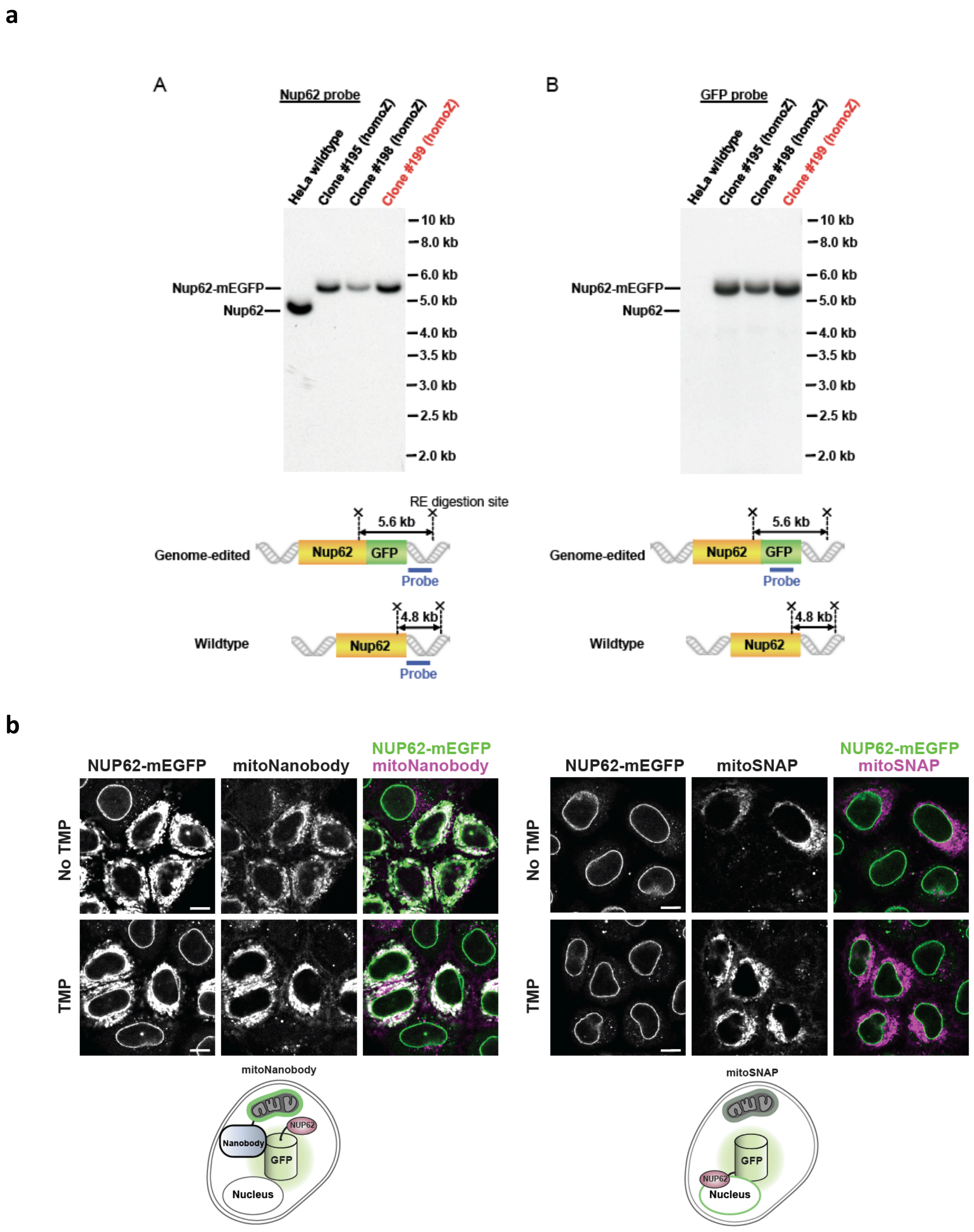

**Supplementary Fig. 13.** Use of $^{GFP}$LAMAs to mislocalize NUP62-mEGFP. (**a**) Homozygousity of NUP62-mEGFP genome edited HeLa Kyoto cell lines were examined by Southern blotting, as previously described[1]. The Clone 199 was used for this study. (**b**) NUP62-mEGFP cells transfected with enhancer nanobody only anchored to the outer membrane of the mitochondria (mitoNanobody), or transfected with SNAP-tag anchored to the outer membrane of the mitochondria (mitoSNAP). Labeled with BG-TMR (10 µM) for imaging of SNAP-tag. TMP = 10 µM. Scale bar, 10 µm.

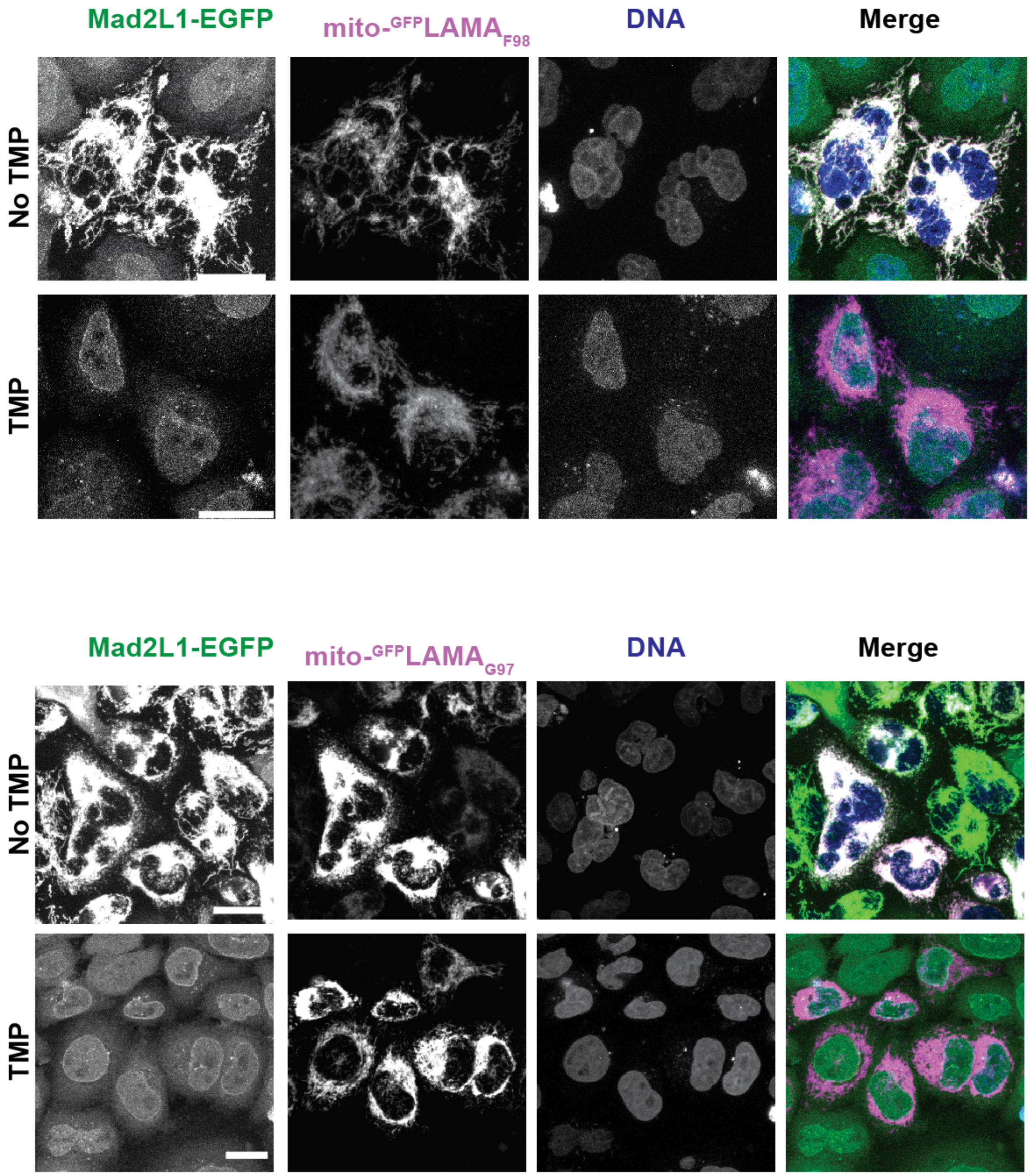

**Supplementary Fig. 14.** Mislocalization of Mad2L1-EGFP using $^{GFP}$LAMAs. Localization of Mad2L1-EGFP in cells transiently transfected with $^{GFP}$LAMAs, in the presence of TMP (10 µM), or after 24 hours washout with media containing DMSO. The LAMAs were labelled with BG-SiR (500nM) to visualize them, and the DNA stained with Hoechst 33342, to score nuclear morphology. Some non-transfected cells are also seen in the image. TMP = 10 µM. Scale bar, 20 µm.

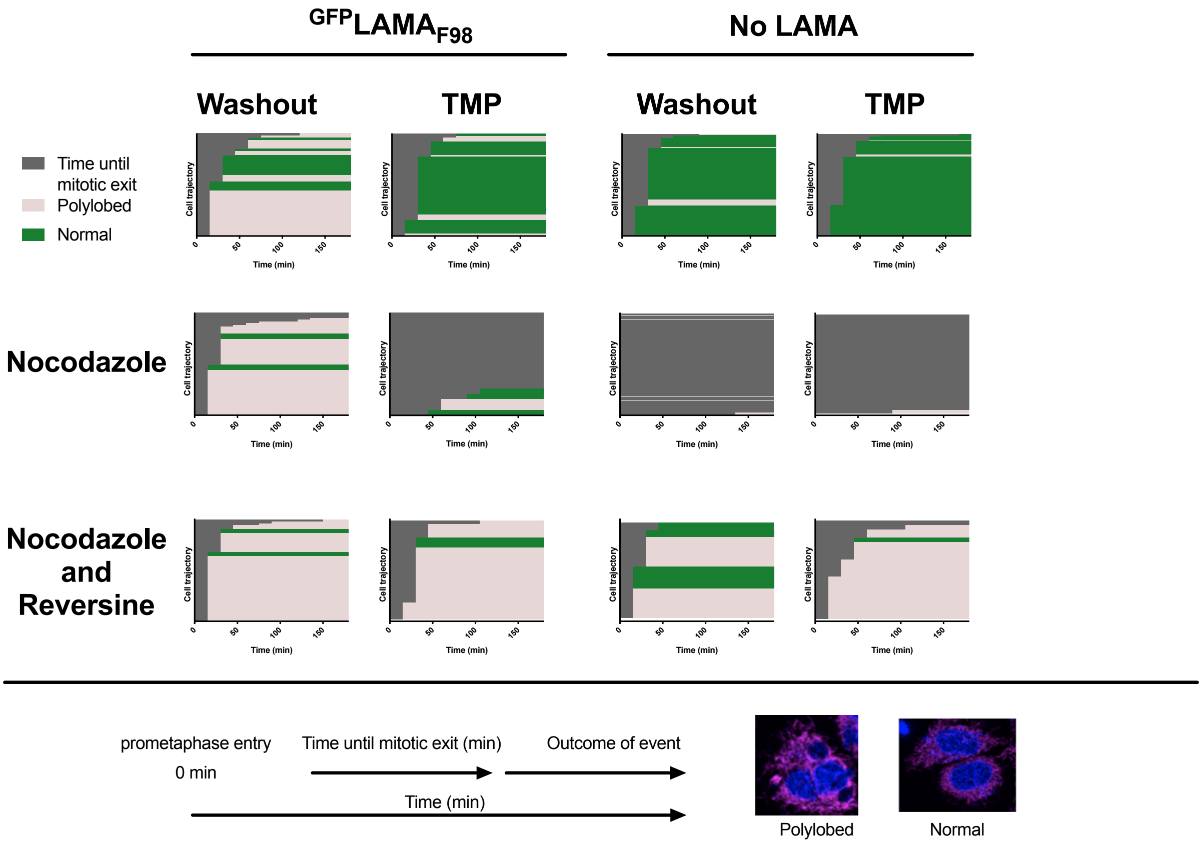

**Supplementary Fig. 15** Cellular trajectories of live-cell imaging of Mad2L1-GFP cells stably expressing $^{GFP}$mitoLAMA$_{F98}$. Cells were imaged every 15 minutes for 20 hours, and aligned at the point at which prometaphase starts (time point 0). Each event was followed until end of mitosis, either by cytokinesis, or decondensation of DNA. After the mitotic event, the nuclear morphology was scored as either polylobed or normal. TMP = 50 µM, nocodazole = 330 nM, reversine = 5 µM. No drug treatment, N = 47, N = 76, N = 79, N =89 cells. Nocodazole treatment: N = 59, N = 19, N = 50, N =27 cells. Nocodazole and reversine treatment: N = 53, N = 29, N = 13, N =23 cells.

a

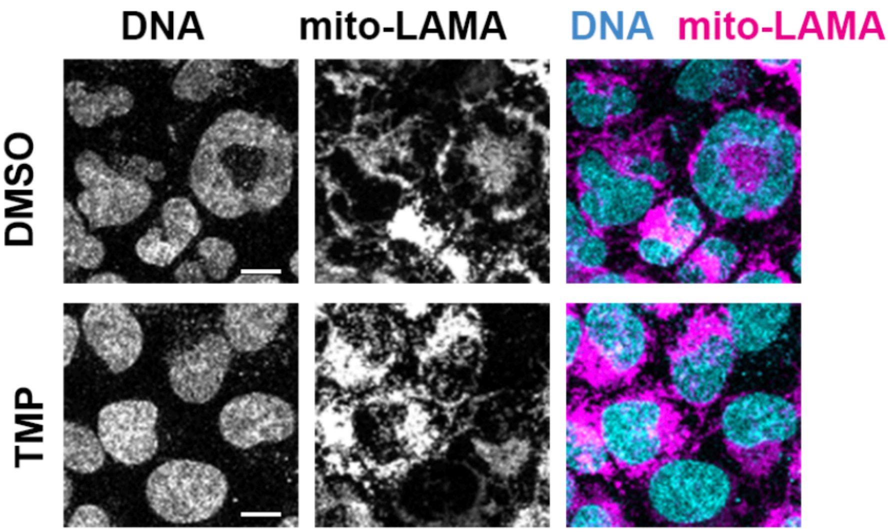

b

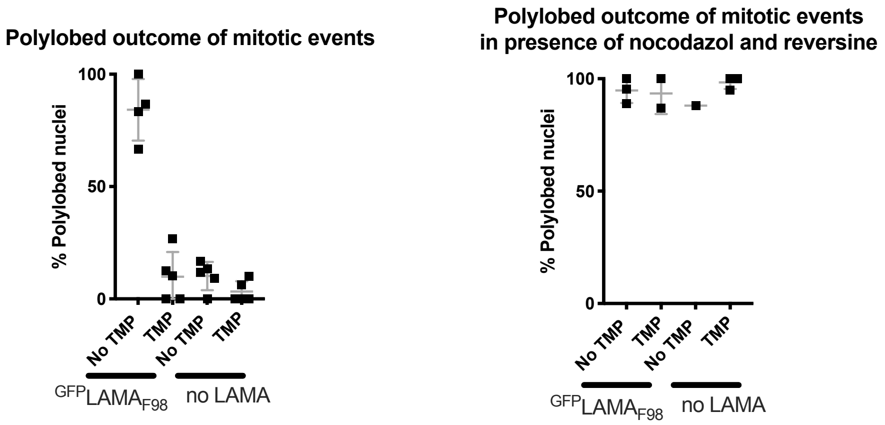

**Supplementary Fig. 16.** Nuclear morphology of Mad2L1-EGFP after mislocalization. (**a**) Nuclear morphology of Mad2L1-EGFP cells stably expressing mito-$^{GFP}$LAMA$_{F98}$ after 20 hours of live-cell imaging after washout of TMP with media containing DMSO, or cells proliferated in TMP. mito-$^{GPF}$LAMA$_{F98}$ was labelled on the SNAP-tag with BG-SiR (100 nM). Scale bar, 10 µm. (**b**) Nuclear morphology manually scored after mitotic events during 20 hours of live-cell imaging. Mean ± s.d. From 3-5 independent experiments with N = 46, N = 75, N = 47, N=92 and N = 52, N = 30, N = 13 and N= 23 cells in each group in total. TMP = 50 µM, nocodazole = 330 nM, reversine = 5 µM.

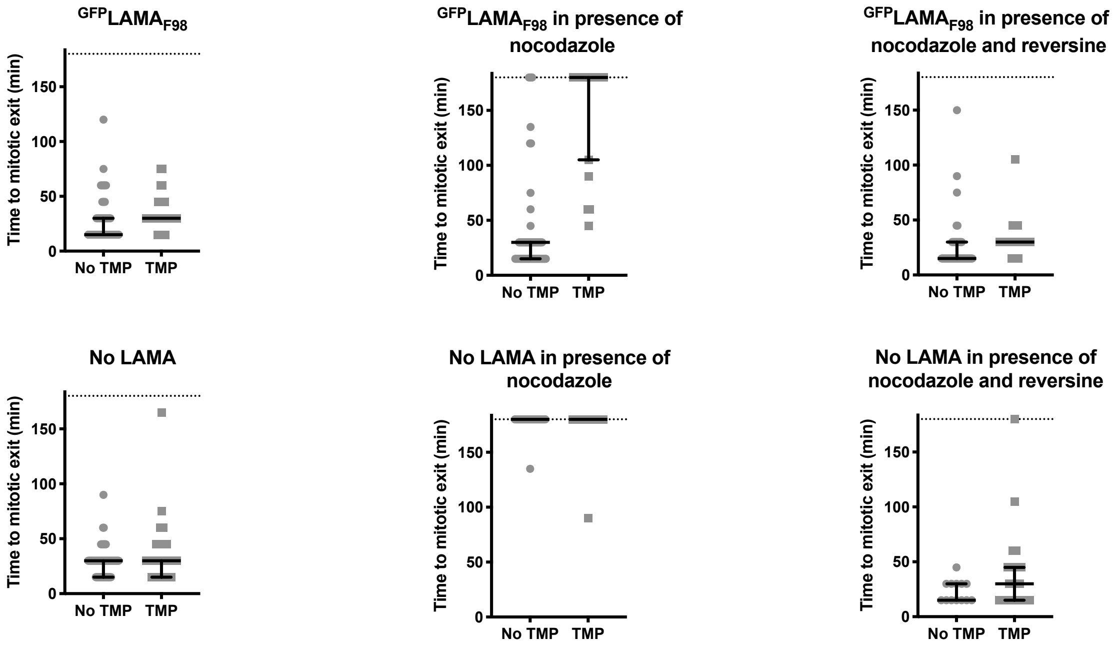

**Supplementary Fig. 17.** Duration of mitosis in HeLa Kyoto Mad2L1-EGFP cells stably expressing mito-$^{GFP}$LAMA$_{F98}$. Time from prometaphase until mitotic exit in Mad2L1-EGFP cells with mito-$^{GFP}$LAMA$_{F98}$, following 20 hours, 15-minute interval, live-cell imaging. Time was cut after 180 minutes (indicated by dotted line). Median ± interquartile range, over 3-5 independent experiments. TMP = 50 µM, nocodazole = 330 nM, reversine = 5 µM.

N = 47, N = 76, N = 63, N = 19, N = 53 and N = 29 for mito-$^{GFP}$LAMA$_{F98.}$ N = 79, N = 92, N = 50, N = 27, N = 13 and N = 23 cells for No LAMA.

**Supplementary Videos 1-4.** Live-cell imaging of Mad2L1-GFP expressing a mito-LAMA. Live cell imaging of a HeLa Kyoto genome edited to Mad2L1-EGFP cell line, and stably expressing mito-$^{GFP}LAMA_{F98}$ (Video 1,2) or no LAMA (Video 3,4). The cells had been grown in the presence of TMP (50 µM), which was washed out at time point 0, or kept during the experiment. The mitochondrially anchored protein constructs were expressed as a SNAP-tag fusion, continuously labeled with BG-SiR (100 nM) to follow expressing cells and for autofocusing purposes. Nuclear morphology was visualized with Hoechst 33342. Transmission images were also recorded to visualize overall cell morphology. Scale bar, 20 µm.

**Supplementary Video 1.**
Genome edited Mad2L1-EGFP cells stably expressing SNAP-tagged mito-$^{GFP}LAMA_{F98}$. TMP (50 µM) was washed out at time point 0.

**Supplementary Video 2.**
Genome edited Mad2L1-EGFP cells stably expressing SNAP-tagged mito-$^{GFP}LAMA_{F98}$ and kept in TMP (50 µM) during imaging.

**Supplementary Video 3.**
Genome edited Mad2L1-EGFP cells stably expressing SNAP-tag on the outer membrane of the mitochondria. TMP (50 µM) was washed out at time point 0.

**Supplementary Video 4.**
Genome edited Mad2L1-EGFP cells stably expressing SNAP-tag on the outer membrane of the mitochondria and kept in TMP (50 µM) during imaging.

**Supplementary Table 1.** X-ray data collection and refinement statistics

| | $^{GFP}$LAMA$_{F98}$ (PDB ID: XXX) | $^{GFP}$LAMA$_{G97}$ (PDB ID: XXX) |
|---|---|---|
| **Data collection** | | |
| Space group | $P6_2$ | $P12_11$ |
| Cell dimensions | | |
| *a, b, c* (Å) | 86.6, 86.6, 73.0 | 47.9, 56.6, 53.3 |
| $\alpha, \beta, \gamma$ (°) | 90.00, 90.00, 120.00 | 90.00, 94.91, 90.00 |
| Resolution (Å) | 50-2.2 (2.3-2.2)* | 50-1.6 (1.7-1.6)* |
| No. unique reflections | 15911 (1970)* | 37239 (6123)* |
| $R_{merge}$ | 0.058 (0.453)* | 0.054 (0.490)* |
| $I/\sigma I$ | 22.6 (5.3)* | 13.2 (2.4)* |
| Completeness (%) | 99.9 (99.9)* | 99.1 (98.6)* |
| Redundancy | 10.1 (10.4)* | 3.3 (3.2)* |
| Wilson B (Å$^2$) | 50.7 | 26.1 |
| **Refinement** | | |
| Molecules per a.u. | 1 | 1 |
| Resolution (Å) | 43.3-2.2 | 47.7-1.6 |
| No. unique reflections | 15910 | 37239 |
| $R_{work}/R_{free}$ | 0.1747/0.2195 | 0.1830/0.2275 |
| No. atoms | 2298 | 2574 |
| Protein | 2167 | 2187 |
| Ligand/ion | 79 | 92 |
| Water | 52 | 295 |
| *B*-factors | 46.3 | 23.1 |
| Protein | 46.5 | 21.8 |
| Ligand/ion | 39.1 | 20.9 |
| Water | 46.6 | 33.8 |
| R.m.s. deviations | | |
| Bond lengths (Å) | 0.008 | 0.006 |
| Bond angles (°) | 1.068 | 0.957 |
| Ramachandran | | |
| Favoured (%) | 98.5 | 99.3 |
| Outliers (%) | 0 | 0 |
| Clashscore | 5.7 | 5.6 |

*Values in parentheses are for highest-resolution shell.

**Amino acid sequences of LAMAs used:**

**Color guide:**

Enhancer nanobody

cpDHFRL24G5

cb9 nanobody

**$^{GFP}$LAMA$_{F98}$**

MAQVQLVESGGALVQPGGSLRLSCAASGFPVNRYSMRWYRQAPGKEREWVAGMSSAGDRSSYEDSV
KGRFTISRDDARNTVYLQMNSLKPEDTAVYYCNVNVGFLPADLAWFKRNTLNKPVIMGRHTWESIGRP
LPGRKNIILSSQPGTDDRVTWVKSVDEAIAACGDVPEIMVIGGGRVYEQFLPKAQKLYLTHIDAEVEGDT
HFPDYEPDDWESVFSEFHDADAQNSHSYCFEILERRGGGGGMISLIAALAVDRVIGMENAMPWNEYW
GQGTQVTVSS

**$^{GFP}$LAMA$_{G97}$**

MAQVQLVESGGALVQPGGSLRLSCAASGFPVNRYSMRWYRQAPGKEREWVAGMSSAGDRSSYEDSV
KGRFTISRDDARNTVYLQMNSLKPEDTAVYYCNVNVGLPADLAWFKRNTLNKPVIMGRHTWESIGRPL
PGRKNIILSSQPGTDDRVTWVKSVDEAIAACGDVPEIMVIGGGRVYEQFLPKAQKLYLTHIDAEVEGDTH
FPDYEPDDWESVFSEFHDADAQNSHSYCFEILERRGGGGGMISLIAALAVDRVIGMENAMPWNFEYW
GQGTQVTVSS

**$^{GFP}$LAMA$_{N95}$**

MAQVQLVESGGALVQPGGSLRLSCAASGFPVNRYSMRWYRQAPGKEREWVAGMSSAGDRSSYEDSV
KGRFTISRDDARNTVYLQMNSLKPEDTAVYYCNVNLPADLAWFKRNTLNKPVIMGRHTWESIGRPLPG
RKNIILSSQPGTDDRVTWVKSVDEAIAACGDVPEIMVIGGGRVYEQFLPKAQKLYLTHIDAEVEGDTHFP
DYEPDDWESVFSEFHDADAQNSHSYCFEILERRGGGGGMISLIAALAVDRVIGMENAMPWNVGFEYW
GQGTQVTVSS

**$^{P24}$LAMA$_{S98}$**

MAQVQLVESGGGLVQAGGSLRLSCAASGSFFMSNVMAWYRQAPGKARELIAAIRGGDMSTVYDDSV
KGRFTITRDDDKNILYLQMNDLKPEDTAMYYCKASGSSLPADLAWFKRNTLNKPVIMGRHTWESIGRPL
PGRKNIILSSQPGTDDRVTWVKSVDEAIAACGDVPEIMVIGGGRVYEQFLPKAQKLYLTHIDAEVEGDTH
FPDYEPDDWESVFSEFHDADAQNSHSYCFEILERRGGGGGMISLIAALAVDRVIGMENAMPWNWGQ
GTQVTVSS

**Additional notes, equations and probe sequences.**

**Fitting models for *in vitro* characterization**

The full equation of single site binding, accounting for the effect of nonspecific binding was used to fit wtGFP fluorescence emission assay and TR-FRET assay:

$$y = (F_{max} - F_{min}) \times Fsb + F_{max} + N \times x$$

where

$$Fsb = \frac{(L + x + K_{\mathrm{d}} - \sqrt{(L + x + K_{\mathrm{d}})^2 - 4 \times L \times x}}{2 \times L}$$

Where $y$ is the emission at 535 nm. $F_{max}$ and $F_{min}$ are the maximum and minimum emission values, and $x$ is the concentration of nanobody protein titrated into wtGFP or the Tb-labeled target. $N$ is a parameter for nonspecific binding, and $L$ is the total concentration of wtGFP or Tb-labeled target ($L$ = 10 nM). $K_{\mathrm{d}}$ is the calculated dissociation constant between the nanobody derivative and wtGFP or Tb-labeled target.

**One phase association and dissociation for kinetic measurements *in vitro* and *in cellulo***

**association**

$$y = NS + (y_0 - NS) \times (1 - e^{-kt})$$

$$t_{1/2} = \frac{\ln(2)}{k}$$

where $y$ is the measured intensity, $y_0$ is the intensity at time 0, $NS$ is the minimal intensity measured at infinite time, $k$ is the pseudo-first-order rate constant, $t$ is the time, and $t_{1/2}$ is the half-time.

**dissociation**

$$y = y_0 + (y_{max} - y_0) \times (1 - e^{-kt})$$

where $y$ is the measured intensity, $y_0$ is the intensity at time 0, $y_{max}$ is the maximal intensity measured, $k$ is the pseudo-first-order rate constant and $t$ is the time.

**ITC**

Proteins were dialysed using mini dialysis kits (GE Healthcare) containing NADPH (1 mM) and/or TMP (0.5 mM/DMSO) as indicated. The protein concentrations were measured by $A_{280}$ using a nanodrop. A Microcal PEAQ-ITC microcalorimeter was used for the measurements. Protein concentration ranged from 10 µM to 40 µM in the cell, and a 10-fold excess in the syringe. 8 or 13-injection programs were chosen depending on the expected affinity between the molecular entities.

**Genome Edited NUP62-EGFP**

The probe sequences for southern blotting are as follows:

**Nup62-mEGFP**

5’AACTTAGTGGCACCAGAGTAACTCTAGTCAGTTACAGTAAAATCCACTGTGTGTGGAAGGCAGAA
GCTAGCGGTTGTATCCCAAGCATCTTTTGTATTTGTCTTTATACTTTGCTGAATTCTCTGAAATACCTA
TTACTGTATGTTGCTTTTCTAAATAAATGTATTGTGAAACCAAAACAGCTGCTGTTAATATGGATAAA
TGTTAGGAGGAGAAAGCTGAGTAAAAAGAGCAGGTTCCAGGAGACTCTGCAGGGGTGCCATTCAC
ATGAAACGCACAGGCAAGCAAATGAAGTAGTGCTTGCATAGACATAGGGGTATGCGATGAAGCAG
CTTTTGTTTGATGAGACAGAGTAATAGACAAATGCAAATCGTGGTTTGCTCCAGGAA3’;

**mEGFP**

5’CACATGAAGCAGCACGACTTCTTCAAGTCCGCCATGCCCGAAGGCTACGTCCAGGAGCGCACCAT
CTTCTTCAAGGACGACGGCAACTACAAGACCCGCGCCGAGGTGAAGTTCGAGGGCGACACCCTGGT
GAACCGCATCGAGCTGAAGGGCATCGACTTCAAGGAGGACGGCAACATCCTGGGGCACAAGCTGG
AGTACAACTACAACAGCCACAACGTCTATATCATGGCCGACAAGCAGAAGAACGGCATCAAGGTGA
ACTTCAAGATCCGCCACAACATCGAGGACGGCAGCGTGCAGCTCGCCGACCACTACCAGCAGAACA
CCC3’.
